# Coupling Metastasis to pH-Sensing GPR68 Using a Novel Small Molecule Inhibitor

**DOI:** 10.1101/612549

**Authors:** Charles H. Williams, Leif R. Neitzel, Pratap Karki, Brittany D. Keyser, Timothy E. Thayer, Quinn S. Wells, James A Perry, Anna A. Birukova, Konstantin G. Birukov, Charles C. Hong

## Abstract

An acidic milieu is a hallmark of the glycolytic metabolism that occurs in cancerous cells. The acidic environment is known to promote cancer progression, but the underlying signaling and cell biological underpinnings of these phenomena are not well understood. Here, we describe ogremorphin, a first-in-class small-molecule inhibitor of GPR68, an extracellular proton-sensing and mechanosensing G protein–coupled receptor. Ogremorphin was discovered in a chemical genetic zebrafish screen for its ability to perturb neural crest development, which shares basic cell behaviors of migration and invasion with cancer metastasis. Ogremorphin also inhibited migration and invasive behavior of neural crest–derived human melanoma cells *in vitro*. Furthermore, in phenome-wide association studies (PheWAS), we identified an aberrantly activated variant of GPR68, which is associated with cancer metastasis *in vivo* and promotes invasive phenotypes of cancer cells *in vitro*. Thus, extracellular proton-sensing GPR68 signaling promotes cell migration and invasion during embryonic development and may do likewise in cancer progression.

## Introduction

Highly malignant, invasive, and metastatic cancers have markedly elevated glycolytic activity that produces an oncologically favorable acidotic extracellular environment, a phenomenon called the Warburg effect ^1^. This moderate acidification (~pH 5.8 to 7.0) of the microenvironment is marked by increases in proton efflux activity through proton ATPases and sodium-proton exchangers ^2–5^. Although acidification promotes tumor malignancy, including metabolic reprogramming and invasiveness, the cellular mechanisms that mediate these phenomena are poorly understood ^6,7^. In numerous animal models of solid tumors, small-molecule inhibition of the sodium-hydrogen antiporter 1 (NHE1) inhibits tumor growth and metastasis ^8^. However, the cellular mechanisms triggered by acidification of the tumor environment are still not wholly understood. One potential signaling pathway involves the extracellular proton-sensing G protein–coupled receptors (GPCRs) GPR4, GPR65, and GPR68, which mediate signaling through Gq and other mechanisms, with maximal activation at pH 6.8–7.0^9,10^. Recently, GPR68 has also been recognized as a mechanosensitive receptor for flow and stretch^11,12^. As this receptor class has been implicated in variety of diseases, including cancer, immunological, pulmonary, and vascular disorders, it is thought to be a promising therapeutic target; however, no small molecule inhibitor of this receptor class has been described^7,13–17^.

During embryonic development, neural crest cells display many cellular behaviors associated with cancer progression: they delaminate and migrate away from a primary site, the neural tube, and disseminate throughout the embryo (migration and invasion) to form diverse cell types and structures such as the peripheral nervous system, smooth muscle cells of cardiovascular system, pigment cells, and craniofacial tissues. In this study, we identified a class of small molecule inhibitors of the proton-sensing G protein–coupled receptor (GPCR) GPR68 that inhibited the migratory behavior of neural crest cells in the zebrafish embryo. This compound class, which we term ogremorphins, also inhibited the migration of cancer cells *in vitro*. In addition, we identified a natural variant of GPR68 associated with metastatic cancers in phenome-wide association studies (PheWAS) and show that aberrant activation of GPR68 promotes invasive phenotypes of cancer cells *in vitro*. Our study suggests that extracellular proton-sensing GPR68 signaling promotes cell migration and invasion during embryonic development and in cancer progression.

## Results

### Discovery of a highly specific GPR68 inhibitor

To discover novel small-molecule modulators of embryonic development, we conducted an unbiased screen of ~30,000 small molecules for their ability to induce phenotypic changes in morphology^18,19^. We identified 5-ethyl-5’-naphthalen-1-ylspiro[1H-indole-3,2’-3H-1,3,4-thiadiazole]-2-one, herein called ogremorphin-1 (OGM1) (Figure 1a) based on its ability to induce abnormal pigmentation (Figure 1b,c). In addition to pigmentation defects, OGM1 reproducibly induced ventral curvature, wavy notochord, shortened body axis, craniofacial defects, and loss of retinal iridophores (Figure 2e). The loss of melanophores, iridophores, and craniofacial cartilage are consistent with defects in neural crest development ^20^. To examine whether the formation of the neural crest was perturbed by OGM1, we stained embryos for *foxD3*, an early marker of the premigratory neural crest, but found no differences between treated and untreated embryos (Supplementary Figure 1). However, time course analysis showed that pigmentation was subject to perturbation by OGM1 during a specific developmental window that correlated to neural crest migration (Figure 1d)^21^. Taken together, these results indicate that OGM1 disrupts development of neural crest–derived tissues subsequent to the formation of neural crest progenitors, most likely during neural crest migration.

**Figure 1.**
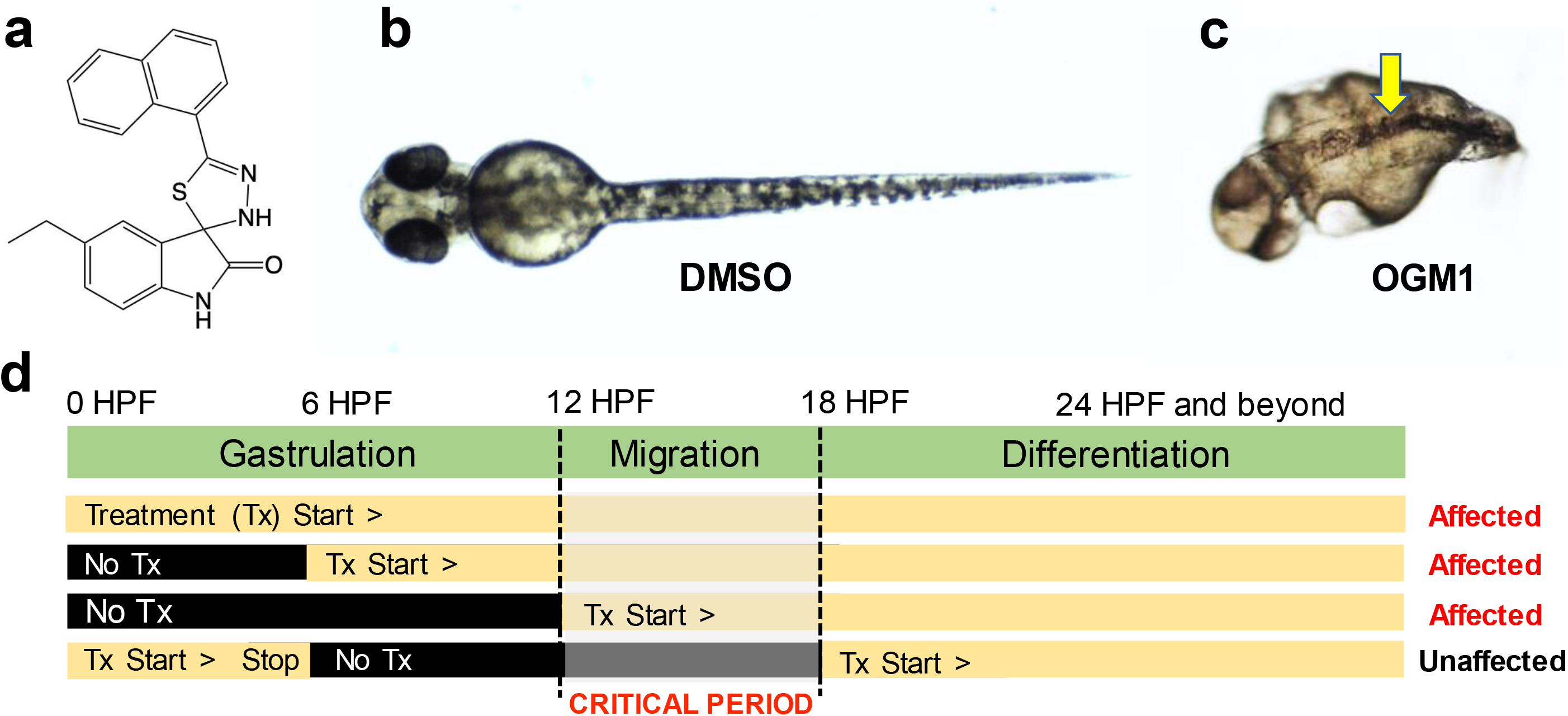
Discovery of OGM1. (**a)** Structure of OGM1, 5-ethyl-5’ -naphthalen-1 -ylspiro[1H-indole −3,2’-3H-1,3,4-thiadiazole]-2-one. (**b)** Dorsal view of DMSO (vehicle control)-treated (CTL) zebrafish embryo at 48 h post-fertilization (hpf). (**c)** Dorsal view of zebrafish embryo treated with 10 μM OGM1 at 48 hpf, showing abnormal pigmentation, characterized by a striped pattern (arrow) restricted to the dorsal aspects of embryo. (**d)** Temporal phenotypic analysis; Embryos treated with OGM1 throughout development (from 0 to 36 hpf) had perturbed pigmentation. Embryos treated from 6 to 36 hpf or from 12 to 36 hpf also had perturbed pigmentation. In contrast, embryos treated from 18 to 36 hpf had normal pigmentation. Embryos treated with a short pulse of OGM1 from 0 to 6 hpf also had normal pigmentation. DMSO = dimethylsulfoxide; Tx, treatment.

**Figure 2.**
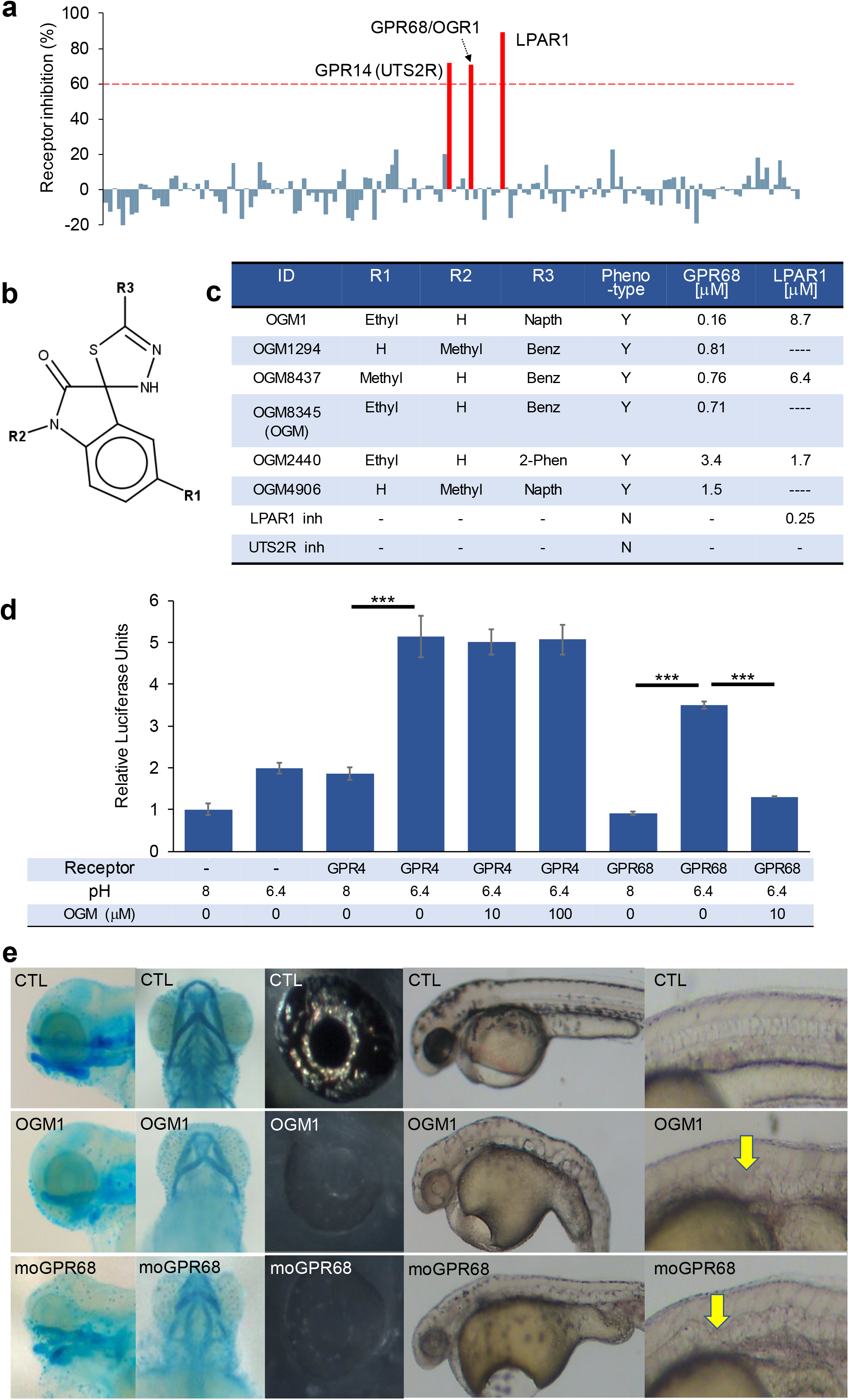
The Ogremorphin-induced phenotype is due to specific inhibition of GPR68. **(a)** Results of Millipore G protein-coupled receptor (GPCR) screen in which 158 GPCRs were tested for agonist and antagonist activity at 10 μM OGM1. Only GPR68/OGR1, LPAR1, and GPR14 (UTS2R) showed significant activity. (**b)** Core scaffold for OGM derivatives. (**c)** Chemical linkage analysis. Loss of LPA activity did not correlate with loss of phenotype in zebrafish. Commercial inhibitors (inh) of UTS2R (SB657510, Sigma) and LPAR1 (Ki16425, Sigma) did not recapitulate the phenotype. **(d)** Representative results from serum-responsive element (SRE-) luciferase assays. SRE-luc had low basal levels of activity in 293T cells. Upon cotransfection with GPR4, luciferase activity increased with acidification but was not inhibited by OGM8345 (OGM) at 1, 10, and 100 μM. In comparison, when GPR68 was cotransfected with SRE and stimulated by acidification, OGM completely inhibited the signal at 10 μM. n = 4, \*\*\**P* <0.001. **(e)** Morpholino knockdown of GPR68 led to craniofacial dysmorphogenesis, loss of iridophores, disrupted pigmentation and a wavy notochord, all of which were also seen with Ogremorphin treatment (Phenotypes quantified in Supplementary Figure 5). CTL, control. MoGPR68 = GPR68 morpholino treatment.

As GPCRs represent >30% of targets for FDA-approved small molecules and are known to modulate cell migration, we assessed OGM1’s activity against 158 GPCRs in a single point assay using Chem-1 cells that use a promiscuous G protein to trigger calcium flux. We found that OGM1 significantly inhibited only LPAR1 (Lysophosphatidic Acid Receptor 1), GPR14/UTS2R (Urotensin-II Receptor), and GPR68/OGR1 (Ovarian Cancer G-coupled Protein Receptor 1) (Figure 2a). Chemical segregation analysis was conducted to further identify the inhibitory activity responsible for the zebrafish phenotype. Commercially available inhibitors of LPAR1 (Ki16425) and GPR14 (SB657510) failed to induce a noticeable phenotype at concentrations up to 200× their respective IC50s of 50 μM and 36 μM, respectively. Next, a small-scale structure-activity relationship (SAR) study around the core pharmacophore of spiro[1H-indole-3,2’-3H-1,3,4-thiadiazole]-2-one generated 3 molecules that were similar to OGM1 but lacked LPAR1 activity (Figure 2b,c). The GPR68 inhibitory activity of spiro[1H-indole-3,2’-3H-1,3,4-thiadiazole]-2-one analogs segregated with the ability of the molecule to induce the zebrafish phenotype; furthermore commercially available LPAR1 and UTS2R inhibitors did not phenocopy the OGM1-induced phenotypes. For further experiments, OGM8345 (henceforth called OGM or Ogremorphin) was resynthesized and used due to its submicromolar potency and GPR68 specificity (Supplementary Figures 2 and 3).

Zebrafish GPR68 and human GPR68 have a high degree of homology, with 58.3% identity and 75.2% similarity (Supplementary Figure 4a). In comparison, the other proton-sensing GPCR family members GPR4 and GPR65 are less similar to GPR68, exhibiting just 41.45% and 28.04% similarity, respectively (Supplemental Figure 4b).

Nevertheless, we wondered whether OGM inhibited GPR4 as well. To test this, we examined the effects of OGM on acid-induced SRE (serum responsive element):luciferase activity in cells transfected with GPR4 or GPR68 and found that OGM inhibited signaling only in cells transfected with GPR68 (Figure 2d). Thus, OGM is a selective inhibitor of GPR68.

To determine if the loss of GPR68 activity was sufficient to cause the phenotypic defects seen in zebrafish treated with OGM (Figures 1c, 2e), we knocked down GPR68 expression using morpholino oligonucleotides. We observed dose-dependent neural crest–specific phenotypes in iridophores, melanocytes, and craniofacial cartilage using 1.5 ng and 3 ng morpholino, whereas the same amount of the mismatch morpholino did not recapitulate these results (Figure 2d, Supplementary Figure 5). In addition, both OGM treatment and GPR68 knockdown caused a wavy notochord (Figure 2e), which is not a known neural crest-related phenotype. Finally, we assayed inhibitors of various proton extrusion mechanisms and found that the H^+^/K^+^ ATPase antagonists omeprazole and lansoprazole generated the same phenotypes as OGM treatment (Supplementary Figure 6). Taken together, these data suggest that the phenotypes observed with OGM treatment are due to the loss of proton-sensing GPR68 activity and include defective neural crest development and wavy notochord formation.

Recently, GPR68 has been described also as a mechanosensor in the endothelium responsible for flow-mediated dysfunction^12^. To test whether OGM or OGM1 also inhibit mechanosensory signaling by GPR68, we used a previously described model of lung endothelial cell dysfunction caused by cyclic stretch of magnitude (CS; 18%) associated with injurious mechanical ventilation with high tidal volumes ^12,22,23^. Our results suggest increased GPR68 mRNA expression in endothelial cells (ECs) undergoing 18% CS (Figure 3a). The GPR68 inhibitors OGM and OGM1 reduced GPR68 mRNA expression to control levels. Previous reports by our and other groups showed magnitude-dependent activation of RhoA signaling in EC exposed to CS ^24,25^. In agreement with published studies, 18% CS activated the Rho pathway, as indicated by pronounced activation of the small GTPase RhoA and increased phosphorylation of myosin phosphatase targeting protein (MYPT) and myosin light chain (MLC) (Figure 3b,c). EC pretreatment with the GPR68 inhibitors OGM or OGM1 attenuated CS-induced activation of Rho signaling (Figure 3b,c). We have previously shown that EC exposure to 18% CS increases expression of inflammatory markers ^22,23^. In agreement with our previous data, stretch preconditioning elevated tumor necrosis factor α (TNFα) mRNA levels (Figure 3d) and increased expression of EC-specific vascular cell adhesion molecule 1 (VCAM-1) (Figure 3e). Remarkably, GPR68 inhibitors suppressed EC inflammatory activation in response to 18% CS for each of these readouts. Taken together, these data suggest that the OGM class of compounds specifically inhibits both the proton-sensing and mechanosensing actions of the GPR68 receptor.

**Figure 3.**
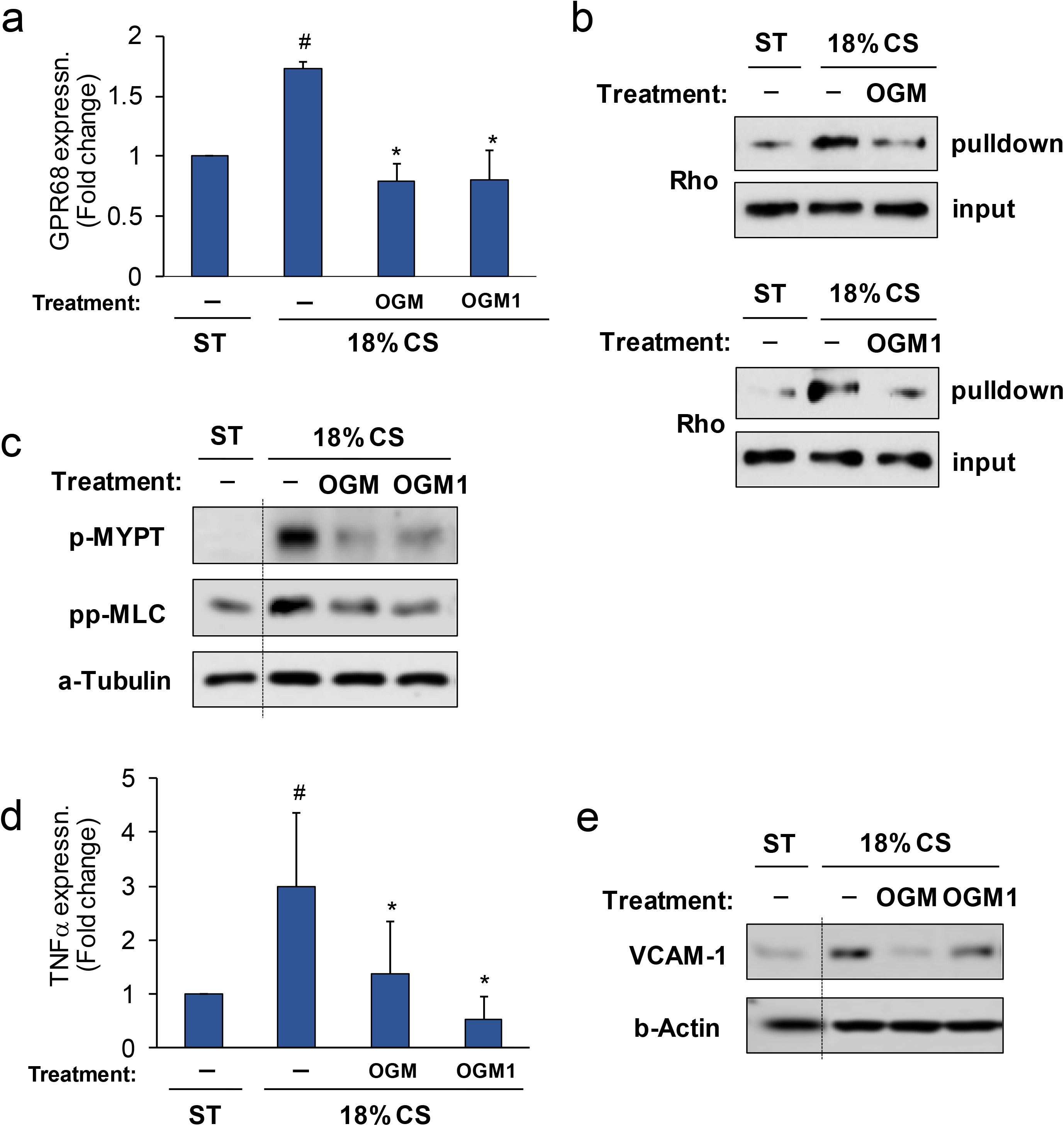
Ogremorphin inhibits cyclic stretch-induced activation of the RhoA pathway and inflammatory signaling. Human pulmonary endothelial cells (ECs) were pretreated with OGMs (OGM at 5 μM and OGM1 at 3 μM) for 30 min. Cells were exposed to either static conditions (ST) or 18% cyclic stretch (CS). **(a)** Cyclic stretch (CS) for 3 h increased *GPR68* mRNA expression in EC. This was by abolished by OGM and OGM1. ^#^*P* < 0.05 vs. ST, \**P* < 0.05 vs. 18% CS alone. Error bars indicate standard deviation. **(b)** CS for 15 min activated Rho in ECs, in a GPR68 dependent manner, as determined by Rho-GTP pulldown assay with or without GPR68 inhibitors (OGM or OGM1). **(c)** CS for 15 min activated myosin phosphatase target (MYPT) and myosin light chain (MLC) in ECs, as determined by their phosphorylation. These were blocked by OGM and GOM1. **(d)** CS for 3 h induced *TNF-α* mRNA expression in ECs. This was by abolished by OGM and OGM1. n = 3, ^#^*P* < 0.05 vs. ST, *P<0.05 vs.18% CS. Error bars, standard deviation. **(e)** CS induced VCAM-1 expression in ECs. This was by abolished by OGM and OGM1.

### Inhibition of GPR68 impairs melanoma migration

As there are numerous parallels between migrating neural crest cells and cancer cells, we examined whether OGM could inhibit the migration of human melanoma cells, which arise from neural crest-derived melanocytes ^26–28^. Previous studies with the MV3 melanoma cell line revealed a pH gradient at the extracellular surface of migratory melanoma cells, with the pH at the leading edge lower than the pH in the rest of the cell and that disruption of proton efflux by inhibition of sodium-hydrogen antiporter 1 (NHE1) with 5-(N-Ethyl-N-isopropyl)amiloride (EIPA) reduced melanoma motility^29–33^. Consistent with these data, in a scratch assay for migration of 3 human melanoma cell lines, A2058, MeWo, and WM115, we found that EIPA reduced migration of A2058 and WM115 cells, although MeWo cell migration was refractory to NHE1 inhibition (Figure 4a,b). OGM significantly reduced migration in all three melanoma lines in the scratch assay compared with vehicle-treated cells (Figure 4a,b). Additionally, OGM completely blocked WM115 cell migration/invasion in the agarose drop assay, indicating that the effect of OGM on cell migration/invasion was particularly pronounced in a dense matrix (Supplementary Figure 7b)^34–36^. To determine which of the proton-sensing receptors are involved in melanoma cell migration, we examined GPR4, GPR65, and GPR68 expression in A2058, MeWo, and WM115 cells. We found that only GPR68 was expressed in all 3 melanoma cell lines and that WM115 cells only expressed GPR68 (Supplementary Figure 7), suggesting that OGM inhibits migration in these cells solely through GPR68 and not the related receptors GPR4 and GPR65. Taken together, these data suggest that proton secretion by NHE1 and proton sensing by GPR68 together regulate melanoma migration *in vitro*.

**Figure 4.**
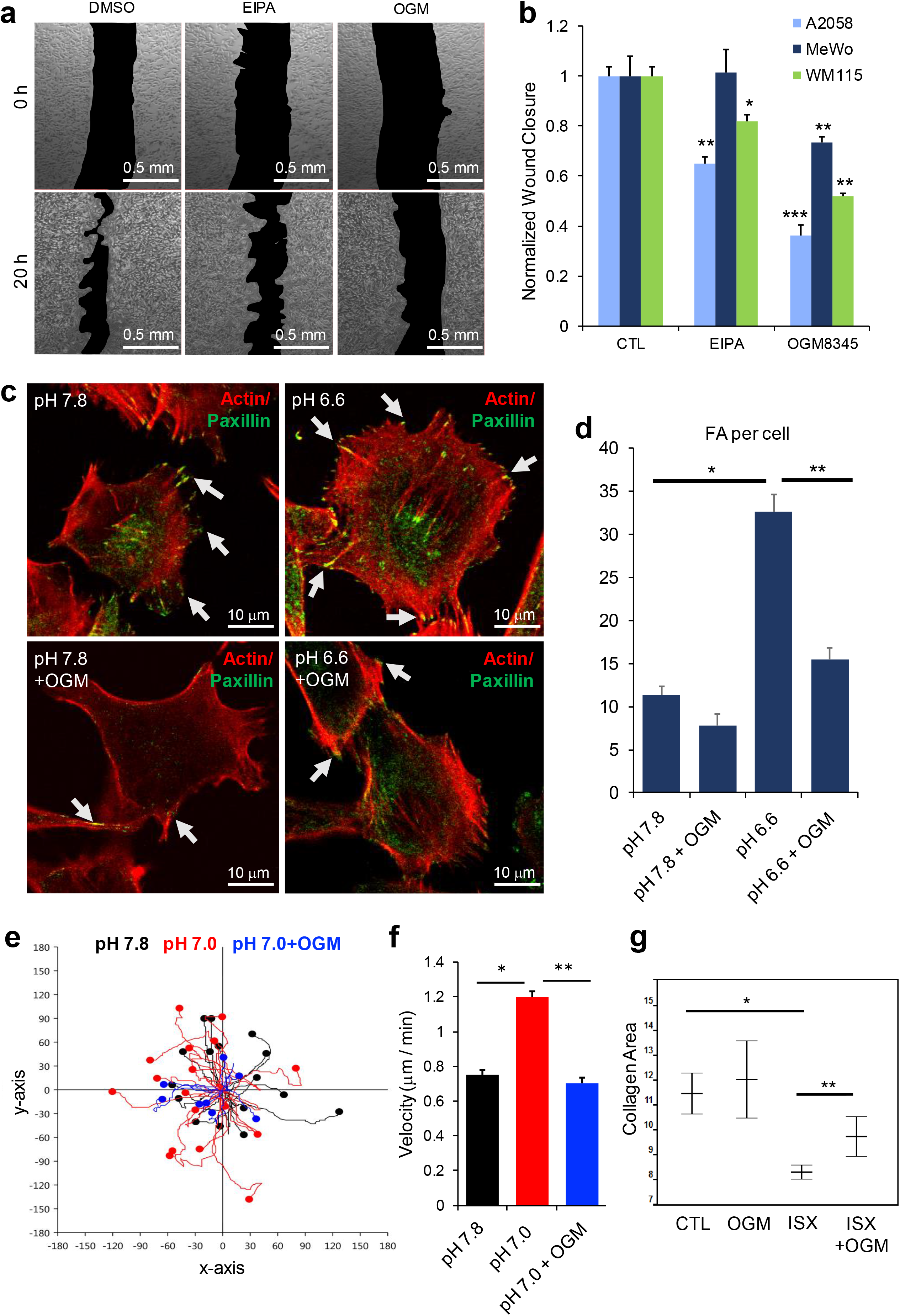
Ogremorphin inhibits human melanoma migration through cell-extracellular matrix interactions. **(a)** In a scratch assay using WM115 cells, OGM and 5-(N-Ethyl-N-isopropyl)amiloride (EIPA) prevented wound closure compared with the DMSO control (CTL). **(b)** Quantification of scratch assay inhibition across three melanoma lines normalized to the DMSO control. OGM inhibited migration in all three cell lines, whereas EIPA inhibited migration in WM115 and A2058, but not in MeWo. n = 3, \**P* < 0.05, \*\**P* < 0.01, \*\*\**P* < 0.001 vs. their respective controls. **(c)** Focal adhesions (FA; green: paxillin) and F-actin stress fibers (red: F-actin) increased in WM115 cells placed on fibronectin substrates in acidic media (pH 6.6). **(d)** Quantification of focal adhesions per cell, n = 25 cells per condition, \*\**P* < 0.001 vs. pH 7.8, and \*\**P* < 0.001 vs. pH 6.6. **(e)** Representative WM115 cell migration tracks after 30-min exposure to pH change with or without OGM. **(f)** Average WM115 cell velocity during time-lapse. Acidification increased the overall velocity (\**P* < 0.01 vs. pH 7.8), which was inhibited by OGM (\*\**P* < 0.01 vs. pH 7.0). **(h)** Collagen gel contraction assay. We plated 1 × 10^6^ cells per well in a collagen mixture that was allowed to polymerize. After 2 days of treatment, collagen disks were released from the side of the well for imaging and the collagen gel area was measured. Isoxazole (ISX) reduced the collagen gel area (\**P* < 0.001 vs. CTL), and OGM inhibited this effect (\*\**P* < 0.05 vs. ISX), n = 5.

### GPR68 modulates adhesion-contraction in melanoma

Focal adhesions are large, dynamic macromolecule complexes that serve as the mechanical link between intracellular actin bundles and the extracellular matrix and are required for cell motility. We sought to determine if GPR68 plays a role in modulating focal adhesions, as previous studies have shown that the proton efflux machinery, NHE-1, localizes to focal adhesion complexes^30,37^. We stained for the focal adhesion protein paxillin, revealing increased focal adhesions in acidified media that were significantly reduced by OGM (Figure 4c,d). Furthermore, actin staining showed that acidification led to formation of stress fiber–like structures (Figure 4c). At pH 7.8, when GPR68 is inactive, OGM did not impede adhesion to fibronectin, whereas in acidified media (pH 6.7), cells adhered to fibronectin-coated slides in an OGM-sensitive manner (Supplementary Figures 8c,d).

Like focal adhesions, filopodia are sites of cell-matrix interactions and express many of the same proteins. In breast cancer cells, extracellular acidification increased both the number and the length of filopodia ^38^. To see if GPR68 affected filopodia, 293T fibrolasts and B16F11 melanoma cell lines were treated with acidified media. The number of filopodia increased In both cell lines and, in the melanoma cells, was accompanied by a significant increase in filopodia length (Supplementary Figures 9a,b,c,h). These changes in filopodia number and length in fibroblast and melanoma cells were inhibited by co-treatment with OGM (Supplementary Figures 9d,e,h). As acidification could potentially activate many cellular acid sensing mechanisms, we stimulated the cells with the GPR68 agonist isoxazole (ISX) in neutral pH 7.4 media ^39^. We found that the small molecule ISX also increased the number of filopodia in both cell lines and was accompanied by a significant increase in filopodia length in the melanoma cells (Supplementary Figures 9f,g,h).

To determine the acute effect of acidification on the migration of Wm115 cells, we performed time-lapse microscopy, and cell tracking (Figure 4e). A 30-min stimulation by acidic medium increased the total migratory capacity of the Wm115 cells by 50%, with a corresponding 50% increase in overall velocity (Figure 4f). This increase in migration and velocity due to acidification was inhibited by OGM treatment (Figure 4f). One way that an increase in adhesion could increase cell velocity is by also increasing the overall contractile force applied during migration. We used collagen gel contraction assays to assess the effect of GPR68 stimulation on the contractile force generated by WM115 cells. As acidification can change the mechanical properties of collagen, we again stimulated GPR68 with ISX, which decreased the size of the collagen gel; this decrease in area was inhibited by co-treatment with OGM (Figure 4g)^40,41^. These data suggest that GPR68 regulates cell motility through focal adhesions to modulate the mechanics of contraction during migration.

### A gain-of-function variant associated with oncogenic phenotypes in humans

We next sought to determine if GPR68 might play a role in either cancer prevalence or progression in humans by examining large real-world datasets that provide human genome-phenotype associations. In the UK Biobank (UKB), a large cohort study involving 502,650 participants in the United Kingdom^42^, we identified a rare GPR68 variant rs61745752 (minor allele frequency, MAF = 0.00099, GnomAD; or 0.00139, UKB) that causes a truncation at amino acid 335 (E336X) (Figure 5a,b). Mutations in the C-terminal tail of GPCRs can have a number of consequences including loss of desensitization, which is mediated through recruitment of beta-arrestin^43^. Beta-arrestin is recruited through the phorphorylation of three residues in the C-terminal tail^44^. Indeed, the E336X variant lacks the predicted phosphorylation sites and the beta-arrestin binding domain (Figure 5b)^45–47^. To test the functional consequence of this C-terminal truncation, we transfected native GFP-tagged (GPR68-GFP) and truncated GFP-tagged GPR68 (E336X-GFP) GPR68 in HEK293T cells. We found that both receptors were localized diffusely along the cell membrane in unstimulated cells. However, upon acid stimulation, the native GPR68-GFP formed intracellular puncta whereas the E336X-GFP variant did not (Supplementary Figures 10a,b). We tested the functional consequence of this variant in a calcium flux assay, which uses Calbryte 5 dye to monitor intracellular calcium as a measure of Gq activation before and during acidic stimulation. The E336X variant had a higher basal level of calcium staining than the native GPR68, although both variants responded with an approximately two-fold increase in intracellular calcium upon stimulation (Figure 5c). Taken together, these data suggest that rs61745752 encodes an aberrantly active (gain-of-function) GPR68 variant.

**Figure 5.**
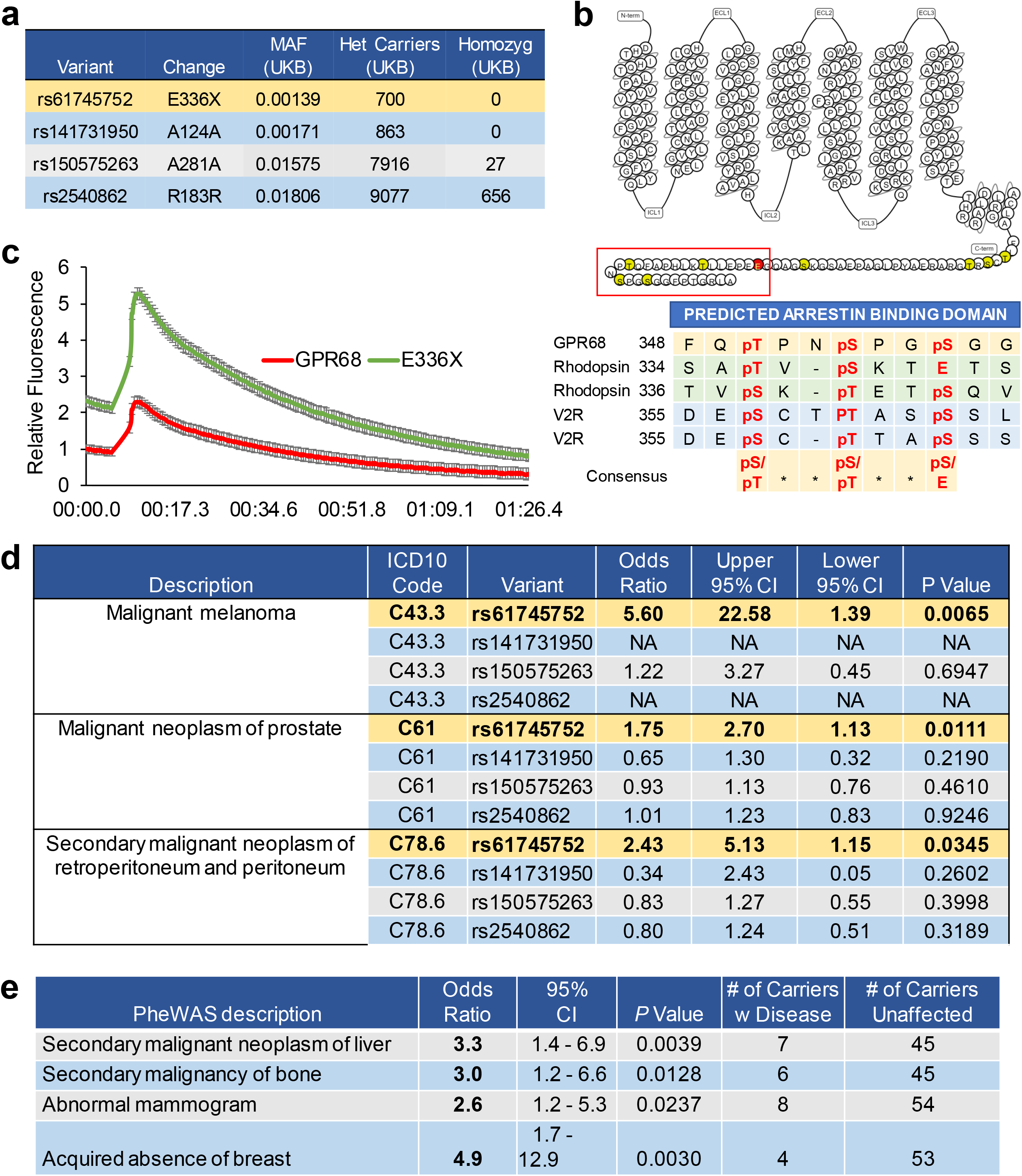
Variant rs61745752 is a gain-of-function allele and correlates with oncological signals in real-world human databases. **(a)** The rare variant rs61745752, which encodes a E336X truncated GPR68, had 700 carriers in UK Biobank. Three synonymous variants of various minor allele frequencies (MAFs) served as negative controls. **(b)** Upper, Snake plot of GPR68 (Red, residue E336, affected by rs61745752; Red box, residues missing in rs61745752; Yellow, putative phosphorylation sites; Lower, conserved beta-arrestin binding motif found in rhodopsin and Vasopressin receptor 2 (V2R) and the putative GPR68 beta-arrestin binding motif lost in rs61745752. **(c)** In calcium flux assays, cell transfected with of the E336X variant had a higher baseline than those transfected with wild-type control, and both respond to acidic stimulation (N = 6 biological replicates). **(d)** In UK Biobank, rs61745752 (E336X) is correlated with cancer-associated ICD10 codes; however, synonymous variants are not. **(e)** In BioVU, rs61745752 is associated with breast cancer-related phenotypes (e.g., abnormal mammogram, acquired absence of breast, mammary dysplasia) and metastasis to liver and bone. Numbers of rs61745752 carriers with and without cancer associated diagnoses.

In the UK Biobank, the rs61745742 variant was associated with the following cancer-related ICD10 codes as either primary or secondary diagnoses (Figure 5d, Supplementary Figure 11a): malignant melanoma (odds ratio [OR]: 5.6; 95% confidence interval [CI]: 1.39-22.58); malignant neoplasm of prostate (OR: 1.75; 95% CI:1.13–2.7); and secondary malignant neoplasm of retroperitoneum and peritoneum (OR: 2.43; 95% CI: 1.15–5.13); (Figure 5d). By contrast, 3 synonymous variants (rs141731950, rs150575263, rs2540862), analyzed as controls (see Methods), were not associated with these or other cancer-related ICD10 codes (Figures 5a,d).

We further investigated the cancer-related diagnoses for the rs61745752 variant in a second cohort of BioVU, a large genomic database linked to Vanderbilt University Medical Center electronic health records ^48–50^. A focused phenome-wide association study (PheWAS) of ~70,000 genotyped participants for association with cancer phenocodes (Supplementary Figure 11b, c; see Methods) showed that this variant was associated with four cancer associated phedoces: secondary malignant neoplasm of the liver (OR: 3.3; 95% CI: 1.4–6.9), secondary malignancy of bone (OR: 3.0; 95% CI: 1.2–6.6), abnormal mammogram (OR: 2.6; 95% CI: 1.2–5.3), and “acquired absence of breast,” which likely indicates mastectomy (OR: 4.9; 95% CI: 1.7–12.9) (Figure 5e) ^48,49^. We manually reviewed the 70 carriers of the rs61745752 variant in the BioVU cohort. Of these, ten had bone or liver metastasis (Supplementary Figure 11d). Notably, seven of the ten were found to have metastatic disease at the time of the initial cancer diagnosis or within 1 year of diagnosis (Supplementary Figure 11d). Focusing on breast cancer, we found that 8 of 70 carriers (11.4%) had a diagnosis of malignant neoplasm of breast, compared with 4.6% in age and sex-matched noncarrier controls (*P* = 0.0136). Moreover, 5.7% (4 of 70) had metastatic breast cancer, versus 1.1% for noncarriers (*P* = 0.0026) (Supplementary Figure 11e). Consistent with these findings, analysis of cBioportal found that patients with breast cancer who harbored mutations in GPR68 had worse overall survival than those without (Supplementary Figure 12a) ^51,52^. By contrast, mutations in GPR4 and GPR65 did not have similar effects on overall survival (Supplementary Figure 12b,c). Taken together with the molecular function of rs61745742, GPR68 could be a clinically relevant target for metastatic potential of various cancer types, including breast cancer.

### GPR68 activity destabilizes cancer spheroids *in vitro*

Next, we sought *in vitro* correlates of metastatic potential using spheroid integrity and invasion assays ^53–57^. Transient transfection of MCF7 breast cancer cells with GFP, GPR68-GFP, and E336X-GFP followed by formation of cell spheroids and subsequent exposure to matrix revealed that a substantial fraction of the spheroids transfected with GPR68-GFP or E336X-GFP displayed disruption of the outer ring of cells by extrusion of inner core cells into the matrix (Figure 6a,b). By contrast, GFP-transfected spheroids maintained their border integrity throughout the duration of the study (Figure 6a,b). Furthermore, the extent of outgrowth of the E336X-GFP-transfected spheroids was significantly greater than that of the GPR68-GFP-transfected spheroids (Figures 6b,c). Notably, for GPR68-GFP- and E336X-GFP-transfected spheroids, GFP-positive cells were found preferentially at the leading edge of cell invasion into the matrix. Conversely, GFP control cells remained evenly spread throughout the spheroid (Figure 6d). Taken together, these data suggest that GPR68 and its E336X variant are sufficient to induce a more invasive phenotype in normally noninvasive MCF7 cells.

**Figure 6.**
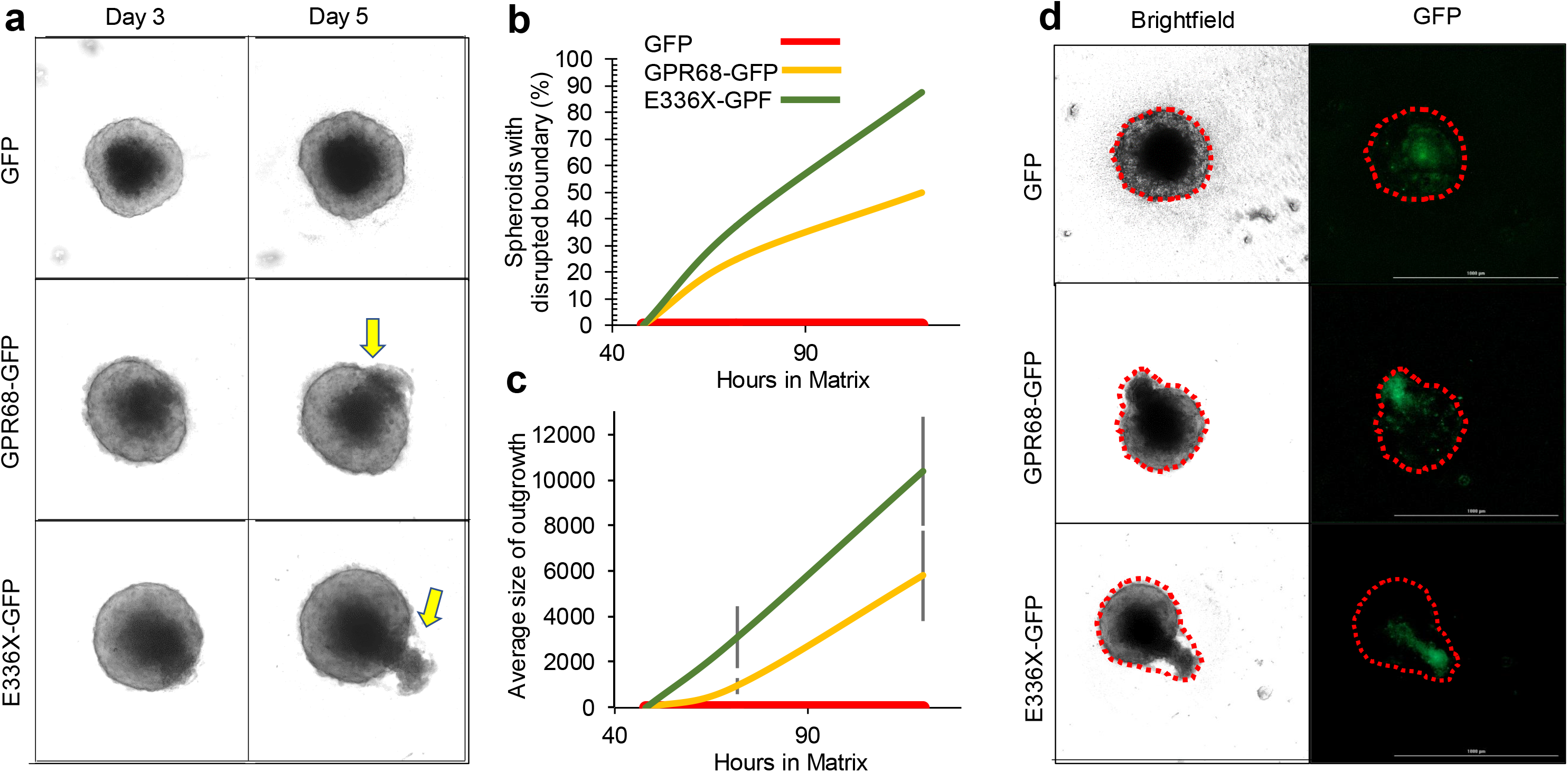
GPR68 activity destabilized MCF7 spheroids. MCF7 cells transfected with GFP, GPR68-GFP, and 336X-GFP were seeded in ultra-low attachment plates to form spheroids. At 48 h, 1:4 galatrox matrix was added. (**a)** GFP-transfected spheroids had the typical structure of an outer proliferative ring with an inner quiescent core. By Day 3, spheroids expressing GPR68-GFP and 336X-GFP had an irregular structure with the outer proliferative ring no longer intact. By day 5, there were outgrowths from the GPR68-GFP and 336X-GFP spheroids that disrupted the outer rings. (**b)** Quantification of the number of spheroids without intact outer rings, N = 8. (**c)** Quantification of the size of the outgrowth from spheroids. Bars: Standard deviation (**d)** Visualization of GFP+ cells in spheroids reveals that they were preferentially located in the spheroid outgrowths. Scale bars = 1,000 μm.

## Discussion

Here, we report the identification of Ogremorphin (OGM), the first-in-class class small molecule inhibitor of GPR68, an unique extracellular proton-sensing and mechanosensing GPCR, based on its ability to disrupt neural crest migration during zebrafish development. OGM inhibited both acid-sensing and mechanosensing GPR68 signaling *in vitro*. Additionally the potent inhibition of cyclic stretch-induced endothelial dysfunction by OGM, suggests an new potential therapeutic target in treatment of vascular diseases associated with perturbed mechanical microenvironment. Mechanistic studies of melanoma with mild acidification (pH 6.5–6.8) induced GPR68 activation resulted in increased melanoma cell migration *in vitro* through regulation of adhesion and contraction. Additionally, we identified a functional gain-of-function variant of GPR68, rs61745752, which encodes a E336X C-terminally truncated receptor and found that it was associated with a variety of malignancies in UK Biobank and BioVU cohorts. Consistent with its association with malignancy in man, an *in vitro* functional assessment of this variant demonstrated that it decreased spheroid integrity and increased invasion. Taken together, our study suggests a role for GPR68 signaling in cancer pathogenesis and a possible therapeutic approach.

A previous study showed that GPR68 is expressed during the early development of zebrafish and is activated by acidification *in vitro* ^58^. Here, we show that inhibition and knockdown of GPR68 disrupt neural crest migration and formation of neural crest– derived tissues, including pigment cells and craniofacial cartilage. As OGM blocks both the mechanosensing and acid-sensing functions of GPR68, we do not formally know if just one or both sensory modalities are involved in neural crest development *in vivo.* In Xenopus, the vacuolar H^+^-ATPase regulates pH in a regional manner to affect craniofacial morphogenesis ^59,60^, and treatment of zebrafish embryos with the H^+^/K^+^ ATPase inhibitors omeprazole or lansoprazole phenocopied OGM treatment. Taken together, these studies suggest that the developmental role of GPR68 we observed is mediated by extracellular proton sensing and imply the existence of acidic extracellular microdomains that regulate cell migration and differentiation during embryonic development. Although such acid microdomains have not yet been demonstrated *in vivo*, an acidified nano-environment has been observed at the leading edges of migrating melanoma cells *in vitro* ^30,31,33^.

Cellular effects of acidification have been studied in numerous cancers ^5–7,30,38,61–66^, and recent studies have revealed various mechanisms by which acidification affects cancer cell behavior through proton-sensing receptors such as GPR4, GPR65, and GPR68^6,67–69^. For example, acidification has been shown to promote migration of melanoma cells in both *in vitro* and *in vivo* models ^32,61,70,71^. Consistent with this observation, our studies with OGM indicate that acid-sensing by GPR68 regulates melanoma cell migratory behavior *in vitro* through regulation of cell adhesion and force generation. Recent studies suggest wide-ranging, sometimes contradictory roles of *GPR68* in cancer progression^17^. For instance, PC3 prostate cancer cells overexpressing *GPR68* have significantly reduced metastatic potential^72^, and HEY1 ovarian cancer cells, overexpressing *GPR68* demonstrate reduced cell migration and increased cell adhesion *in vitro* ^73^. In contrast, GPR68 activation in medulloblastoma enhanced tumor cell migration ^74^. Moreover, GPR68 was found to be a critical regulator of cancer-associated fibroblasts in colon cancer and pancreatic cancer, suggesting an important role of acid-sensing signaling in regulation of the tumor microenvironment ^68,75–77^.

Using a functional genomics approach, we showed that a truncated GPR68 (E336X) variant (rs61745752) missing the C-terminal putative β-arrestin binding domain causes a loss of receptor internalization and an increase in basal calcium levels. Such gain-of-function effects are similar to those observed with the analogous C335X C-terminal truncation of another GPCR, Thyrotropin Releasing Hormone Receptor (TRHR), which results in an ~2× increase in basal Ca+ levels ^78^: We show that the rare rs61745752 variant is weakly assocaited with solid cancers and metastasis in the UK Biobank and with metastatic disease in BioVU. Although specific phecodes did not replicate, the rarity likely precludes replication and a trend of phenotypes have emerged suggesting that GPR68 signaling may be relevant to metastatic disease in humans. The association with metastases was found in diverse, common cancers such as colon, breast, and lung cancer, suggesting the role of GPR68 in the oncogenic processes may be general, rather than cancer-type-specific. Consistent with the association of rs61745742 with human malignancies, we found that MCF7 breast cancer cells expressing the gain-of-function GPR68 variant demonstrate enhanced invasive potential in a tumoroid model that is known to recapitulate the hypoxic and acidic core found in poorly vascularized tumors. Notably, the GPR68-GFP–expressing cells drove the instability by migrating out of the core and pushing out of the periphery of the spheroid. Taken together, our study indicates that GPR68 may be promising therapeutic target that highlights the importance of the acidic microenvironment of solid tumors.

## Acknowledgments

*FoxD3* in situ hybridization probes were a kind gift from the Knapik lab (Vanderbilt University). A2058, MeWo, and WM115 cells were a kind gift from the Quaranta lab (Vanderbilt University). B16F11 were a kind gift from the Tyska lab (Vanderbilt University). MSCV: OGR1-GFP, herein referred to as GPR68-GFP, was a kind gift from Owen Witte (University of California, Los Angeles and Howard Hughes Medical Institute). UK Biobank data were obtained under the UK Biobank Resource Application Number 49852. This work was funded by NIGMS R01GM118557 to CCH.

## Online Methods

### Chemical Screen

All zebrafish experiments were approved by the Vanderbilt University Institutional Animal Care and Use Committee. The chemical screen for small molecules that perturb the dorsoventral axis was performed as described previously ^18,19,79,80^. Briefly, pairs of wild-type (WT) zebrafish were mated, and fertilized eggs were arrayed in 96-well microtiter plates (5 embryos/well) containing 100 μl E3 water. At 4 hpf, a small-molecule library from the Vanderbilt High Throughput Screening Facility was added to each well to the final concentration of 10 μM. Embryos were incubated at 28.5°C and examined for gross morphological changes indicative of embryonic axis dorsalization at 24 and 48 hpf. A total of 30,000 compounds were screened.

### Alcian blue staining

Staged embryos and larvae were anesthetized with tricaine and killed by immersion in 4% formaldehyde (prepared from paraformaldehyde, and buffered to pH 7 in phosphate-buffered saline). Fixed animals were rinsed in acid–alcohol (0.37% hydrochloric acid, 70% ethanol), and stained overnight in Alcian blue (Schilling et al.,1996a). After destaining in several changes of acid–alcohol, preparations were rehydrated. Following rinsing and clearing in a solution of 50% glycerol and 0.25% potassium hydroxide, cartilage was visualized under a stereomicroscope.

### Whole-mount zebrafish in situ hybridization

In situ hybridization was performed as previously described ^81^. Zebrafish foxD3 probes were synthesized as previously described ^82^.

### Target profiling assays for GPCR

GPCR profiling assays were performed by Millipore (St. Louis, MO) in cells expressing Gα15, a promiscuous G protein that enhances GPCR coupling to downstream calcium ion signaling pathways.

### Cyclic stretch

Human pulmonary artery endothelial cells (Lonza, Allendale, NJ) were grown on BioFlex collagen I–coated, 6-well, flexible-bottomed culture plates (Flexcell international Inc., Burlington NC) and cyclic stretch (CS) experiments were performed with an FX-4000T Flexcell Tension Plus system equipped with a 25-mm loading station, as described previously ^22^. Both static and CS cells were seeded into identical density/conditions and stimulations were carried out in the culture medium with 2% FBS. After 48 h in culture, cells were exposed to CS (18% linear elongation, sinusoidal wave, 20 cycles/min.) for the desired period, and static plates were kept in the same incubator. At the end of the experiments, cells were lysed and processed for RNA isolation or western blotting.

### Zebrafish injections

The OGR1 morpholino 5’-TTTTTCCAACCACATGTTCAGAGTC-3’ and the Mismatch morpholino 5’-CCTCTTACCTCAGTTACAATTTATA-3 were synthesized by Gene Tools (Philomath, OR). Morpholinos and mRNAs were injected as described previously^81^.

### Real-time PCR

Total RNA was isolated using the RNeasy Plus Mini Kit (Qiagen, Germantown, MD) following the manufacturer’s instructions. One microgram of RNA was reverse transcribed to cDNA using the iScript cDNA synthesis kit (Bio-Rad, Hercules, CA). The expression levels of different genes were analyzed by qRT-PCR using PerfeCTa SYBR Green Fastmix (Quanta bio, Beverly, MA) on a Bio-Rad CFX96 Real-Time PCR System. Ct values were normalized to GAPDH and fold changes in expression were calculated using the ∆∆Ct method. The following pairs of primers were used for RT-PCR:

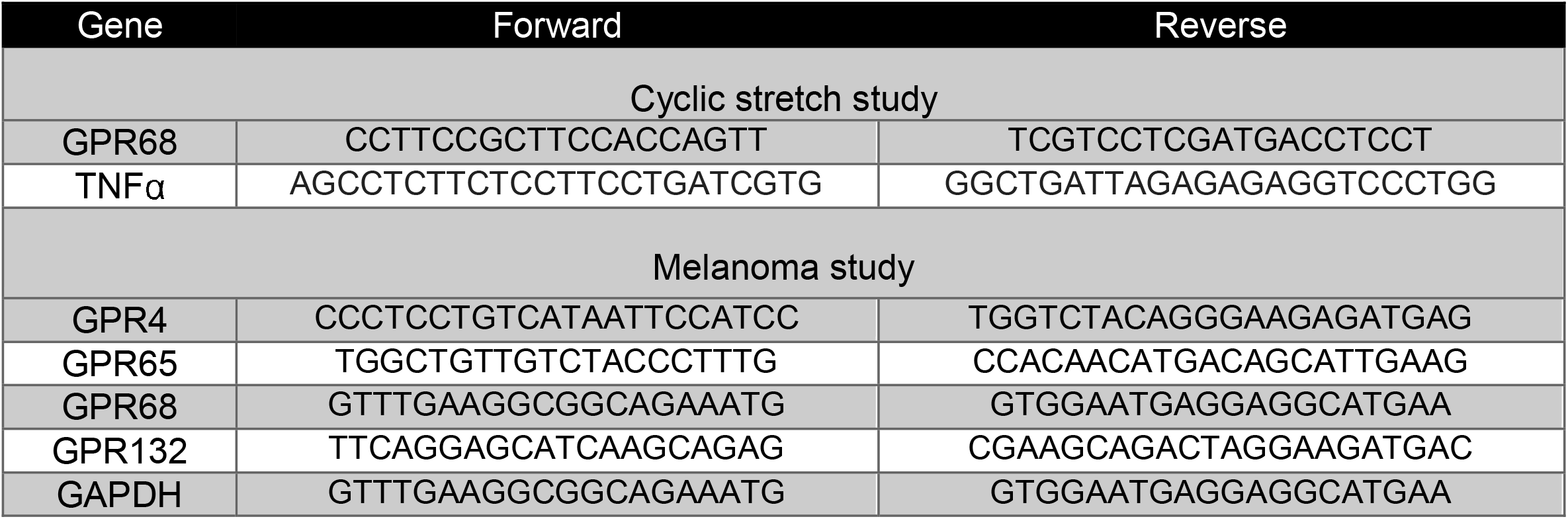

### Cell Culture and Transfection

HEK293, HEK293T, MeWo, A2058, and MCF7 cells were cultured in DMEM supplemented with 10% fetal bovine serum (FBS) (Gibco) and 1% penicillin-streptomycin (Cellgro). WM115 cells were cultured in DMEM/F12 supplemented with 10% FBS and 1% penicillin-streptomycin. Primary human pulmonary artery endothelial cells were obtained from Lonza, Allendale, NJ and propagated in endothelial cell growth medium-2 (EGM-2, Lonza, Allendale, NJ). Cells were transfected with Lipofectamine 3000 according to the manufacturer’s protocols.

### Luciferase transfection and assays

Cells grown in 12-well plates were transfected with 1 μg DNA using Lipofectamine 3000 with sre: luciferase and either GFP, GPR68-GFP, or E336X-GFP. After 3 days, the cells were lysed, and cell extracts were subjected to the Steady-Glo luciferase assay (Promega, Madison, WI), according to the manufacturer’s instructions. The results were normalized to cell titer, as determined using the Cell Titer-Glo luminescence assay (Promega)

### E336X-GFP plasmid generation

A deletion between amino acid 336 and the GFP start codon was generated from the MCSV:GPR68-GFP plasmid (a gift from the Witte lab) using the Q5 mutagenesis kit (New England Biolabs, Ipswich, MA).

### Agarose drop assay

WM115 cells were trypsinized and resuspended at 100,000 cells/ml in low-melting agarose. A 10-μl drop of the cell mixture was added to a 12-well plate. After the agarose solidified, normal culture medium containing either 0.5%DMSO or 5 μM OGM8345 was added. The cells were incubated for 5 days and then the area around the agarose drop was visualized by light microscopy. Cells outside of the agarose drop were quantified.

### Scratch assay

Cells were grown to confluence and then denuded with a P200 micropipette tip. Fresh medium was added and loose cells were removed. The compounds OGM (5 μM) and EIPA (10 μM) were then added. The denuded area was observed 20 h later.

### PheWAS

We performed targeted PheWAS on the single-nucleotide polymorphism (snp) variant in people of european ancestry using methods and code groupings previously described ^49,50^ with the R PheWAS package ^48^. Briefly, comprehensive associations between SNPs and codes for a total of 1,866 clinical phenotypes derived from the International Classification of Disease, 9th Clinical Modification (ICD-9) edition were quantified from the BioVU Cohort’s electronic health records. ICD-9 codes associated with each phenotype can be found at the PheWAS catalog, located online at http://phewas.mc.vanderbilt.edu/. For this targeted analysis we focused on cancer related phecodes. Cases for a given disease were defined as having at least 2 relevant ICD-9 codes on different days. The PheWAS method also defines control groups for each disease, which ensures that related diseases do not serve as controls for the disease currently being analyzed.

### UK Biobank analysis

rs61745752 was identified as potentially deleterious or damaging in multiple pathogenicity scores including DANN, MutationTaster, Eigen scores. The remaining silent mutations were selected based on rarity being comparable to rs61745752 and we extracted the genotype status for all individuals in the UK Biobank at the 4 SNPs in GPR68 and their respective ICD10 codes for neoplasms C00-D49 in primary or secondary diagnoses fields. Carrier versus noncarrier odds ratios were calculated with confidence intervals. As rs61756752 is a very rare variant, we chose to perform a chi-square test of contingency without Yates correction due to its undue strictness for type 1 with smaller sample sizes ^83,84^. All variants were treated the same way for analysis. Additional supplemental analyses were carried out for selected variants and ICD10 phenotypes with a logistic regression model using covariates of age, sex, and 5 principle components.

### Rho GTPase assay

Activation of RhoA GTPase in pulmonary EC culture was analyzed using a GTPase *in vitro* pulldown assay kit available from Millipore (Billerica, MA). Briefly, after desired period of CS exposure, cells were lysed on ice in Mg^++^ lysis/wash buffer (Millipore, Cat no. 20-168) supplemented with 10 µg/ml of leupeptin, 10 µg/ml of aprotinin, 25 mM of NaF, and 1 mM of Na_3_VO_4_. Cell lysates were cleared by spinning at 13000 rpm for 5 min., followed by incubation with pre-washed Rhotekin beads for 1 h at 4 ºC. Beads were washed twice with lysis buffer and beads-bound active Rho proteins were eluted in Laemmli sample buffer by heating at 95 ºC for 5 min. Both active and total Rho proteins were subjected for Western blot analysis.

### Western Blot

Western blot analysis was performed against phospho-myosin phosphatase target (MYPT)-Thr696 (Millipore, ABS45), di-phospho-myosin light chain (MLC) (Cell Signaling, #3674) or Vascular Cell Adhesion Molecule 1 (VCAM-1; Cell Signaling, #136662) antibodies. Probing for α-tubulin (Sigma, T6199) or β-actin (Sigma, A5441) were used as normalization controls.

## Supplementary Figure Legends

**Supplementary Figure 1.**
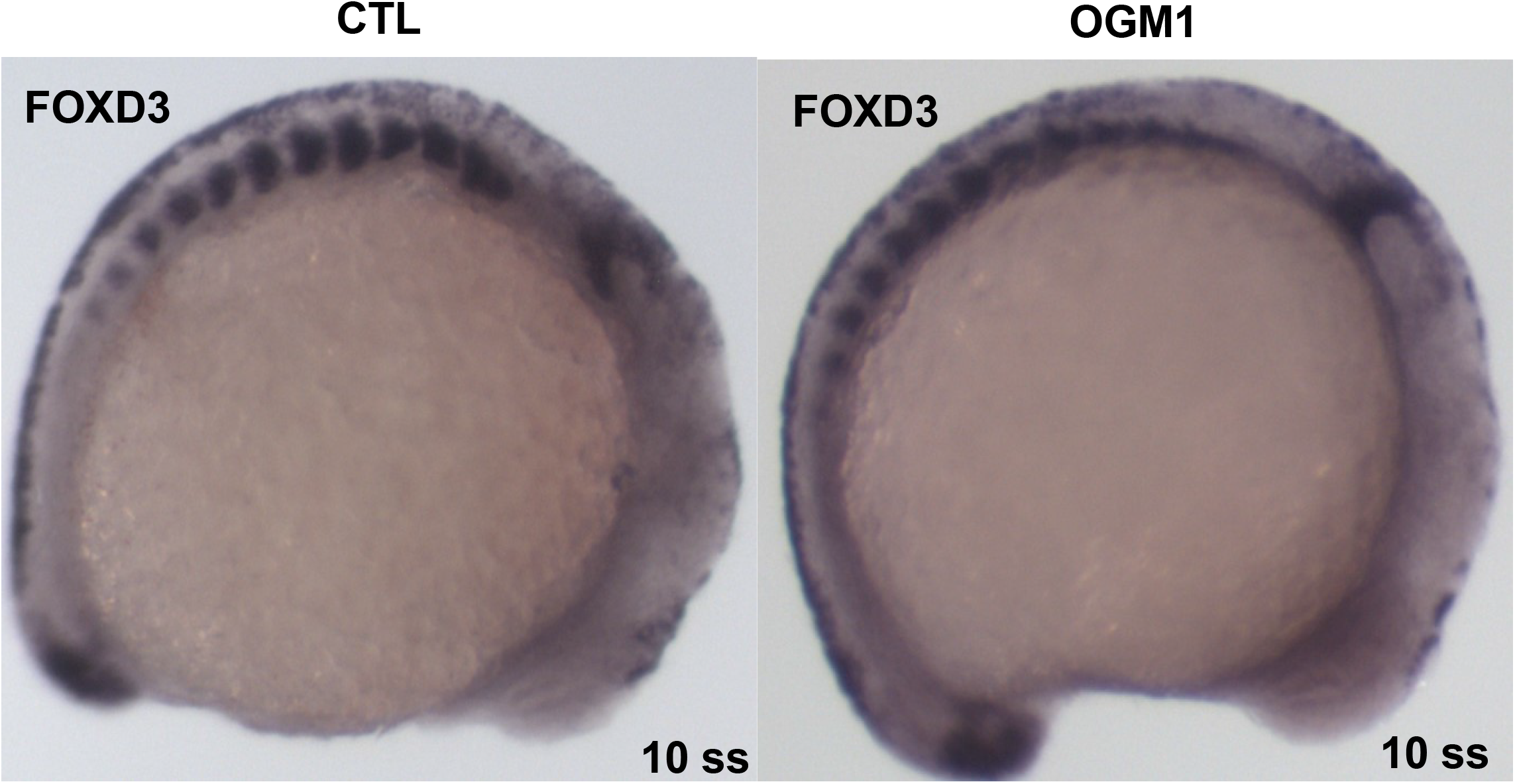
Ogremorphin treatment does not affect neural crest progenitors. Lateral view of zebrafish after *in situ* hybridization of *FOXD3* at the 10-somite stage.

**Supplementary Figure 2.**
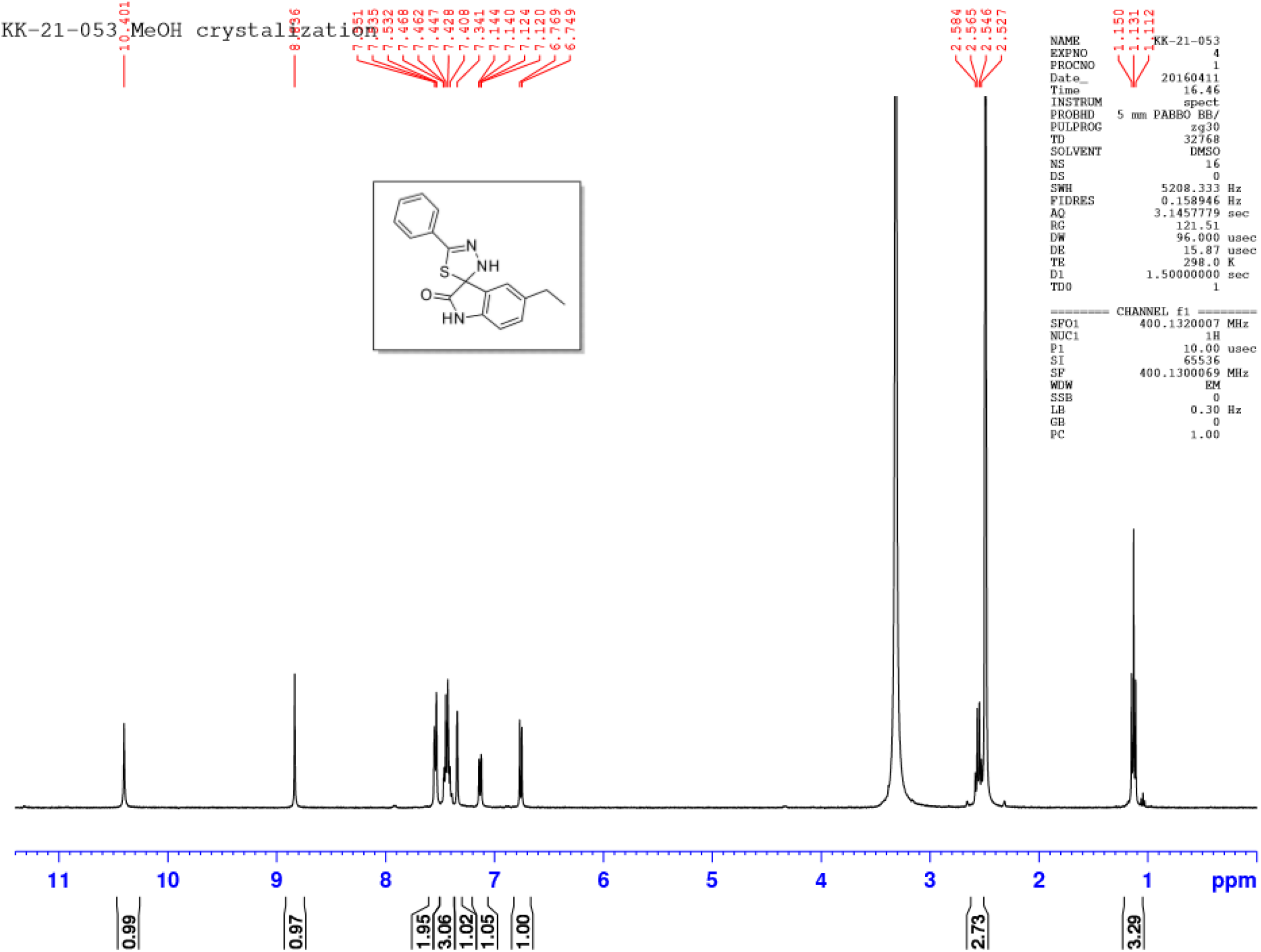
Proton nuclear magnetic resonance of OGM8345 resynthesis.

**Supplementary Figure 3.**
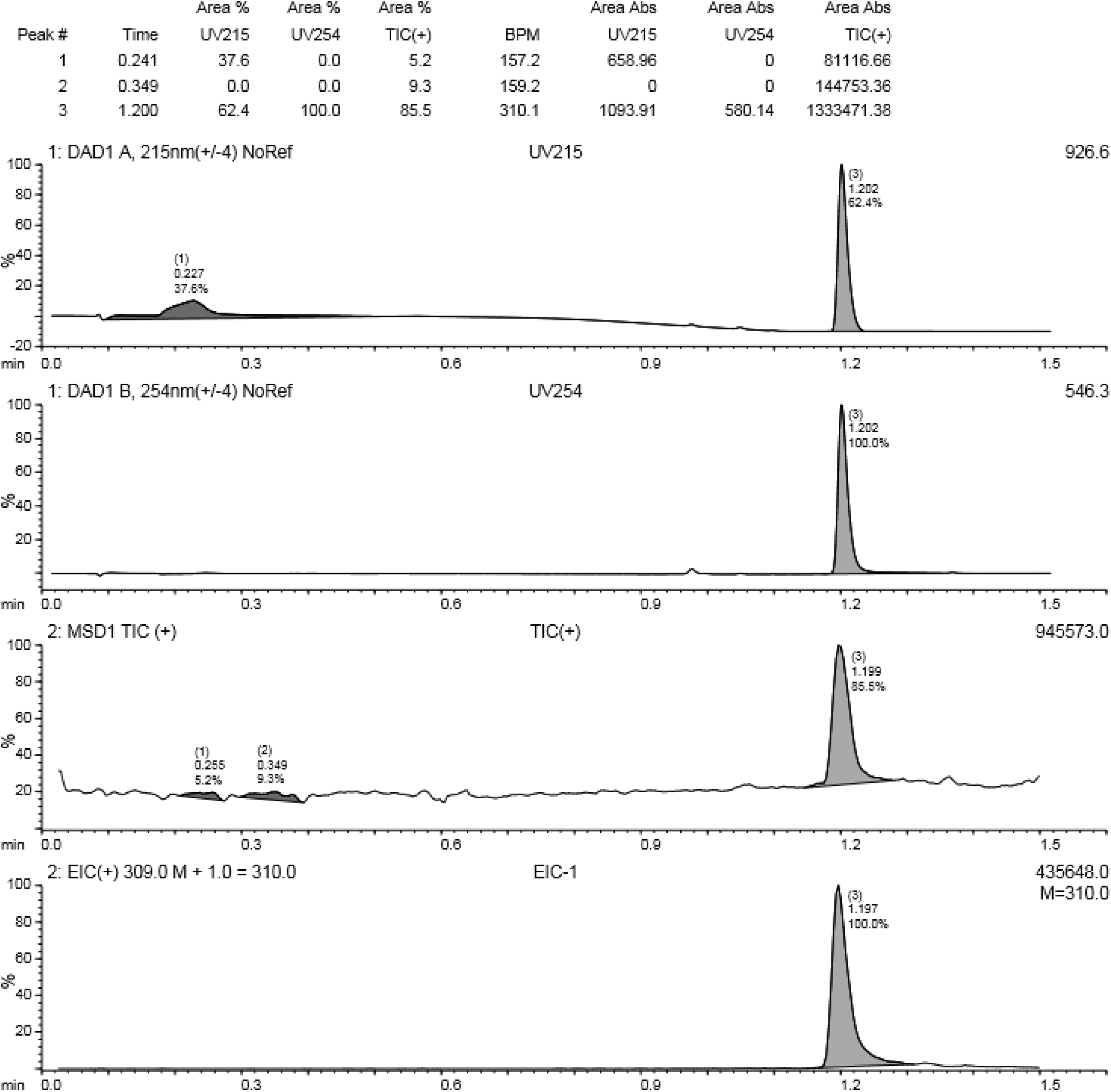
Liquid chromatography/mass spectrometry of OGM8345 resynthesis.

**Supplementary Figure 4.**
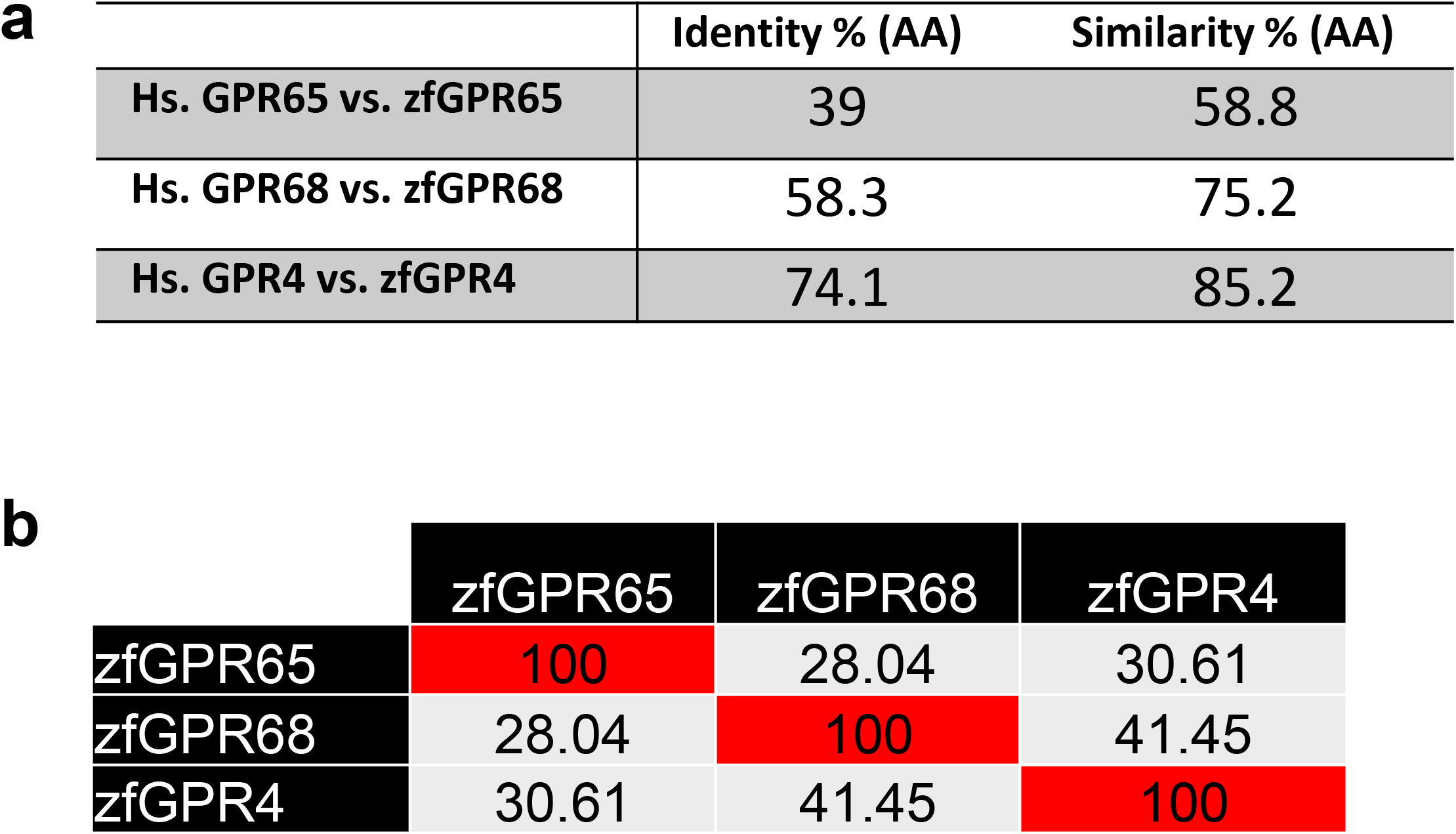
Similarity and Identity between proton-sensing GPCRs. **(a)** Similarity of acid-sensing GPCRs in zebrafish. **(b)** Identity and similarity between human and zebrafish orthologs.

**Supplementary Figure 5.**
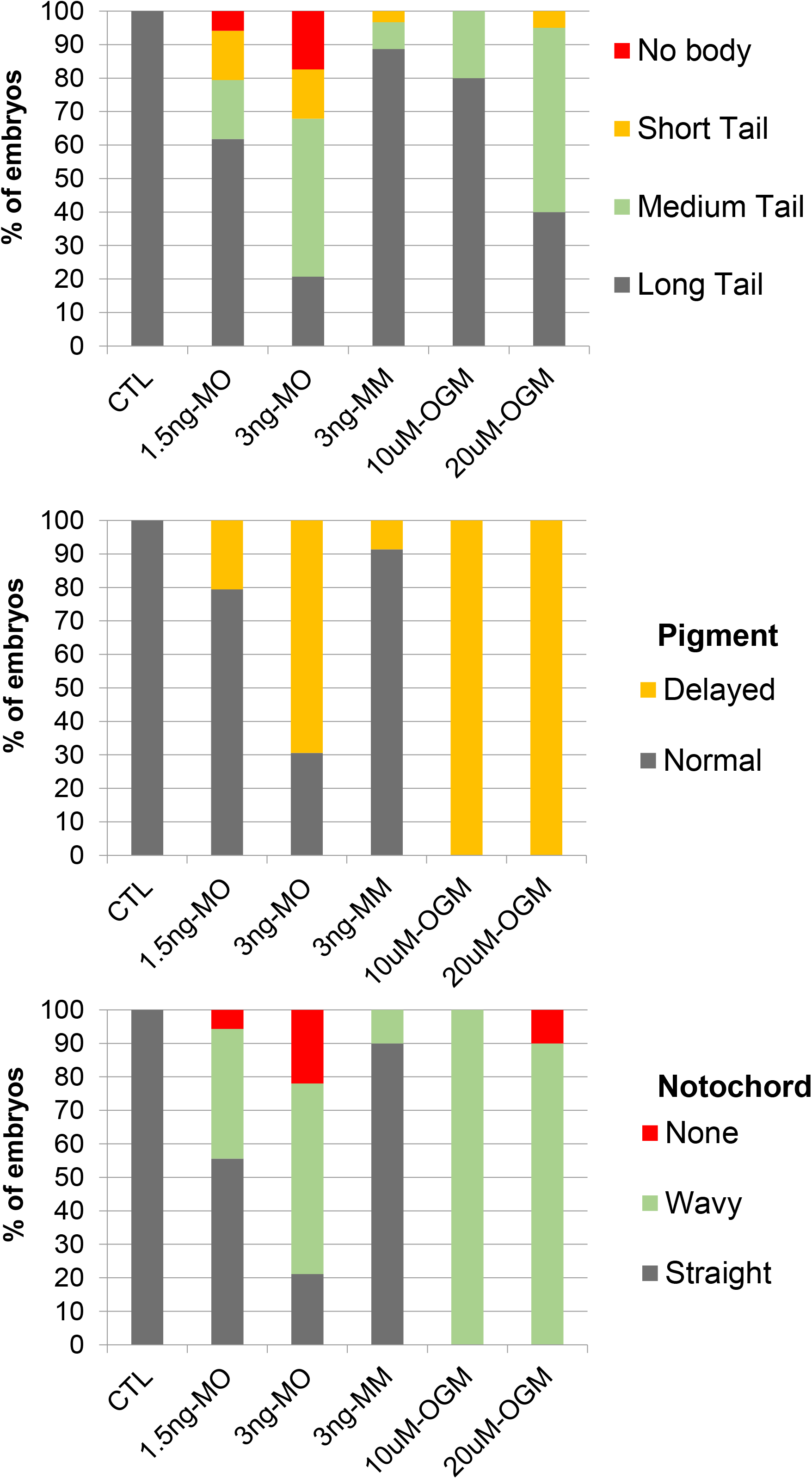
Quantification of GPR68 knockdown and inhibition phenotypes. Morpholino-injected embryos exhibited dose-dependent increases in phenotype prevalence with a shortened body axis, delayed pigmentation, and defects in notochord integrity at 1.5 ng and 3ng. The mismatch morpholino (3 ng) had minimal effects. OGM had increasing effects on these phenotypes at 10 and 20 μM.

**Supplementary Figure 6.**
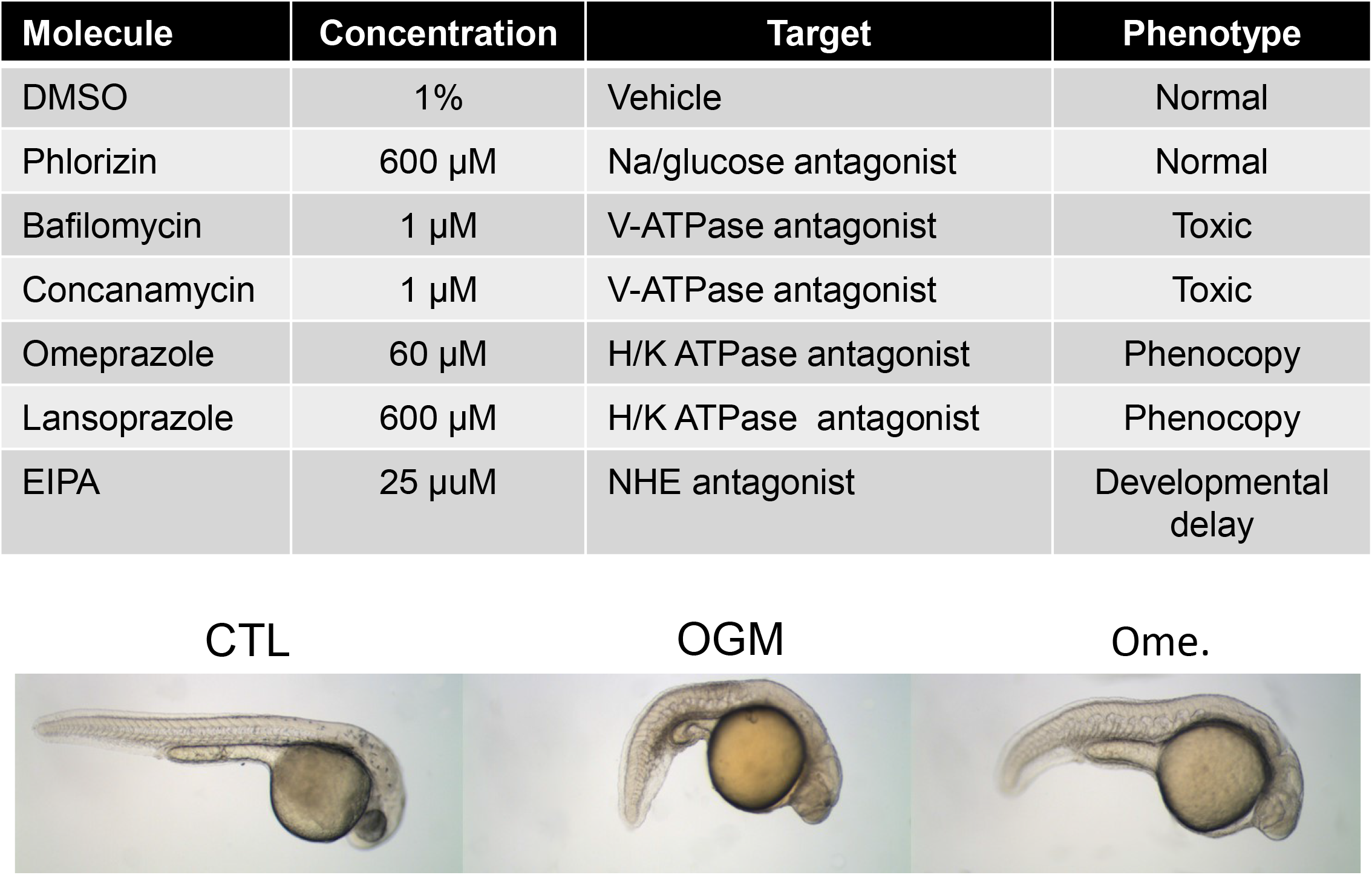
Inhibition of the proton efflux machinery phenocopies OGM treatment. **(a)** Table of proton efflux inhibitors used to treat embryos at 5 hpf. Omeprazole and lansoprazole both phenocopy OGM inhibition. (**b)** Representative images of DMSO-, OGM-, and omeprazole-treated embryos.

**Supplementary Figure 7.**
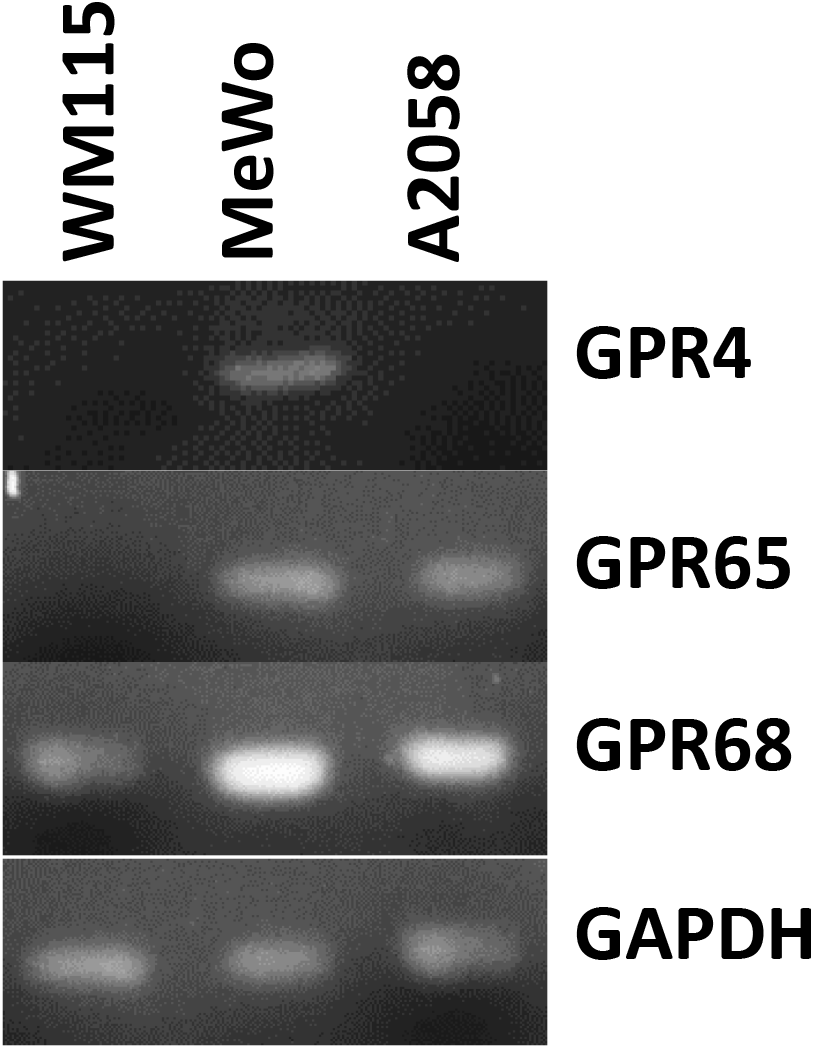
RT PCR of proton-sensing GPCRs in human melanoma cell lines.

**Supplementary Figure 8.**
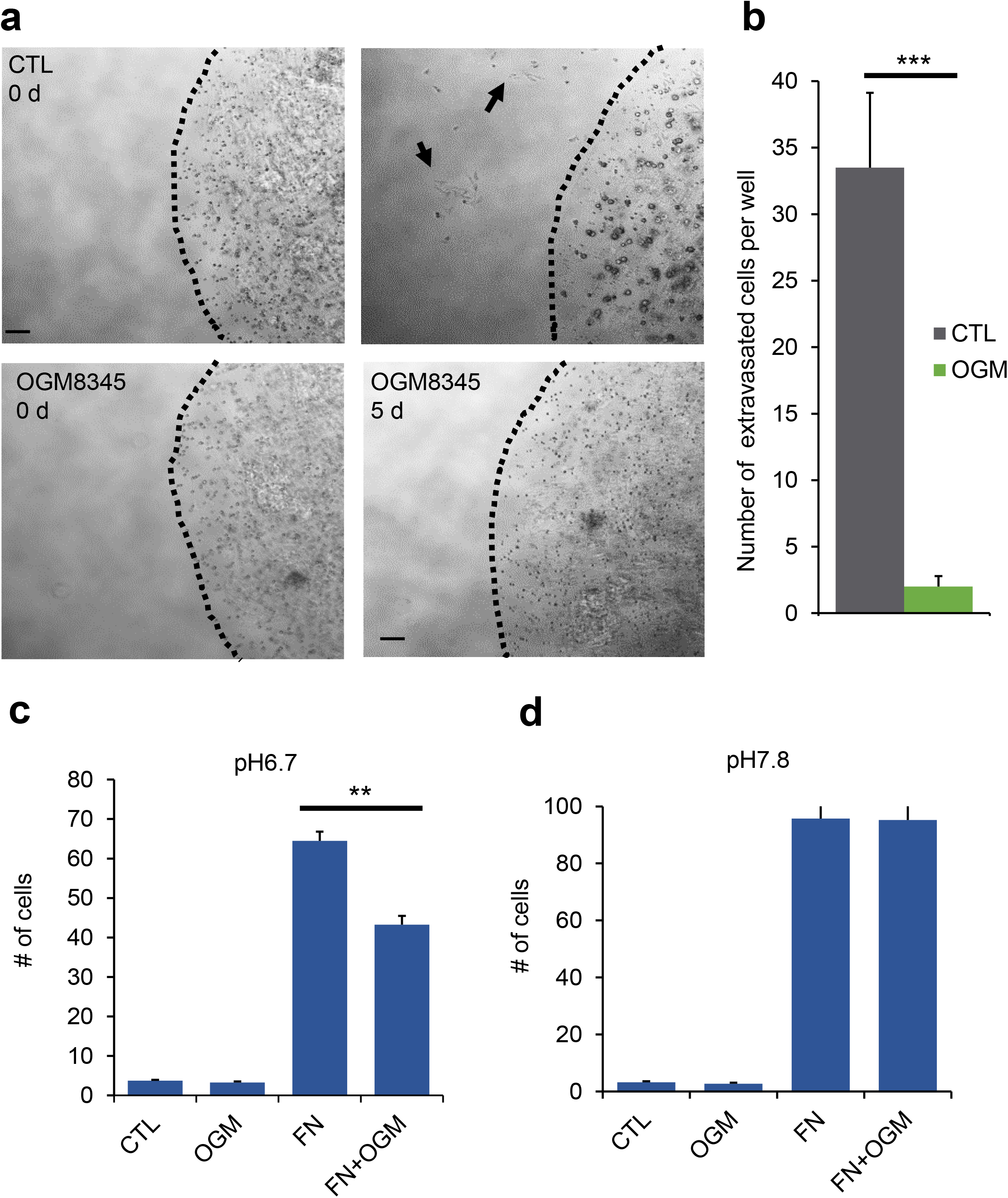
Ogremorphin inhibits migration and adhesion. WM115 cells were mixed into the low melting temperature agarose solution and single 10-μl drops were seeded onto cell culture plates. After the agarose solidified, media containing OGM (5 μM) or control (CTL, 0.05% DMSO) were added, and wells were observed 5 days later. (a) Immediately after addition of media, no cells were on the plate outside of the agarose gel in CTL or OGM wells. Five days after seeding, cells were observed outside of the gel (black arrows) in CTL cells but not in OGM-treated cells. (**b)** Quantification of the number of escaped cells from an average of 5 gel drop replicates. (**c**) Number of WM115 cells adherent to a 96-well plate after 1 h with and without fibronectin coating at pH 6.7. OGM inhibits adhesion (*P* < 0.05). **(d)** Number of WM115 cells adherent to the 96-well plate after 1 h= at pH 7.8. Total number of cells seeded per well = 10,000 for **(c)** and **(d)**.

**Supplementary Figure 9.**
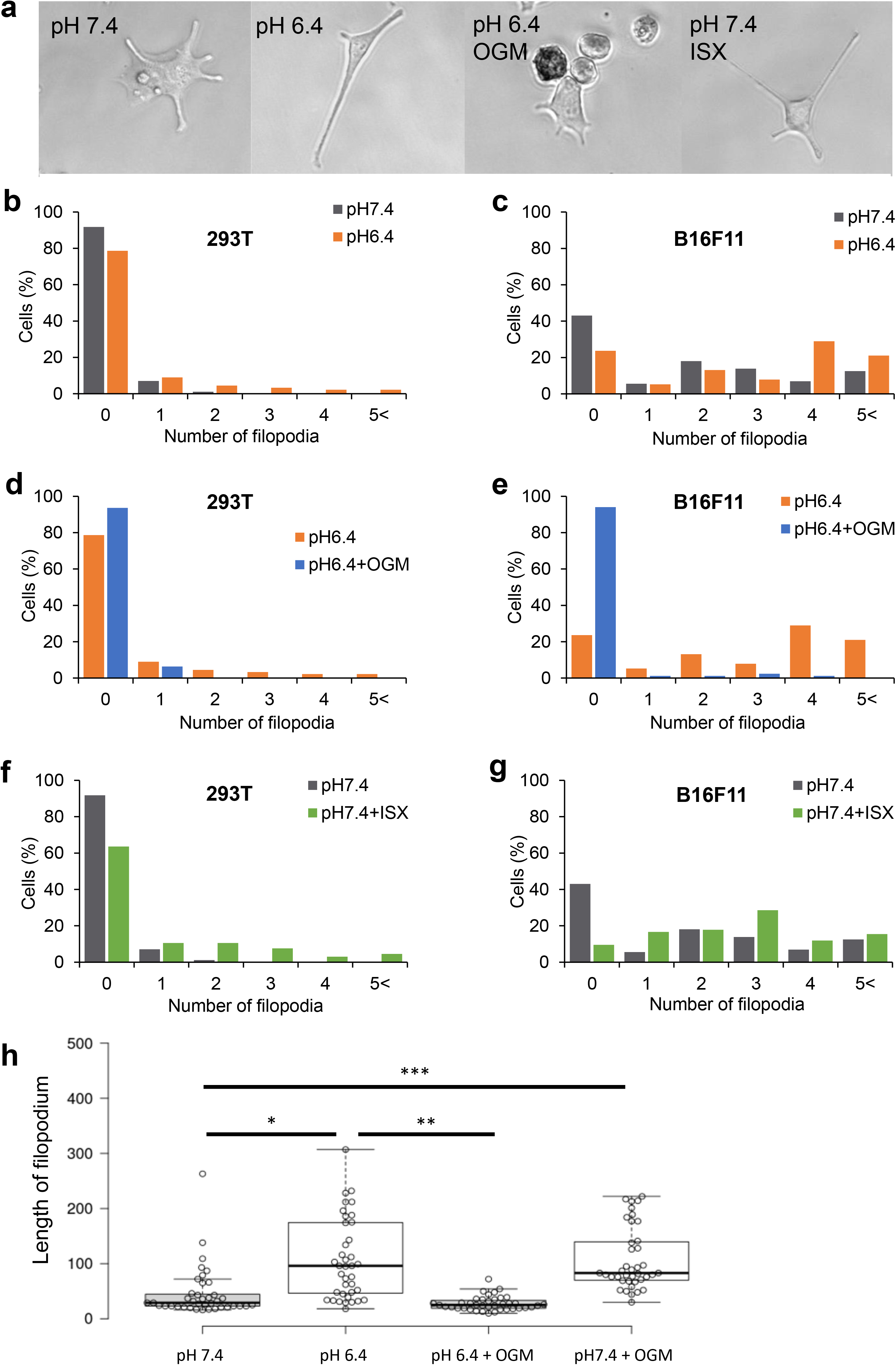
GPR68 stimulation increases filopodial number and length in fibroblasts and melanoma cells. Quantification and comparison of the number of filopodia per cell in 293T cells **(a–c)** and in B16F11 melanoma cells **(d–f)**. Acidification increased the number of filipodia **(a, d)**, whereas OGM treatment decreased the number of filopodia **(b, e).** A GPR68 agonist also increased the number of filopodia per cell in neutral medium **(c, f).** Less than 100 cells from 4 independent experiments were measured per cell line and pH using differential interference contrast microscopy imaging.

**Supplementary Figure 10.**
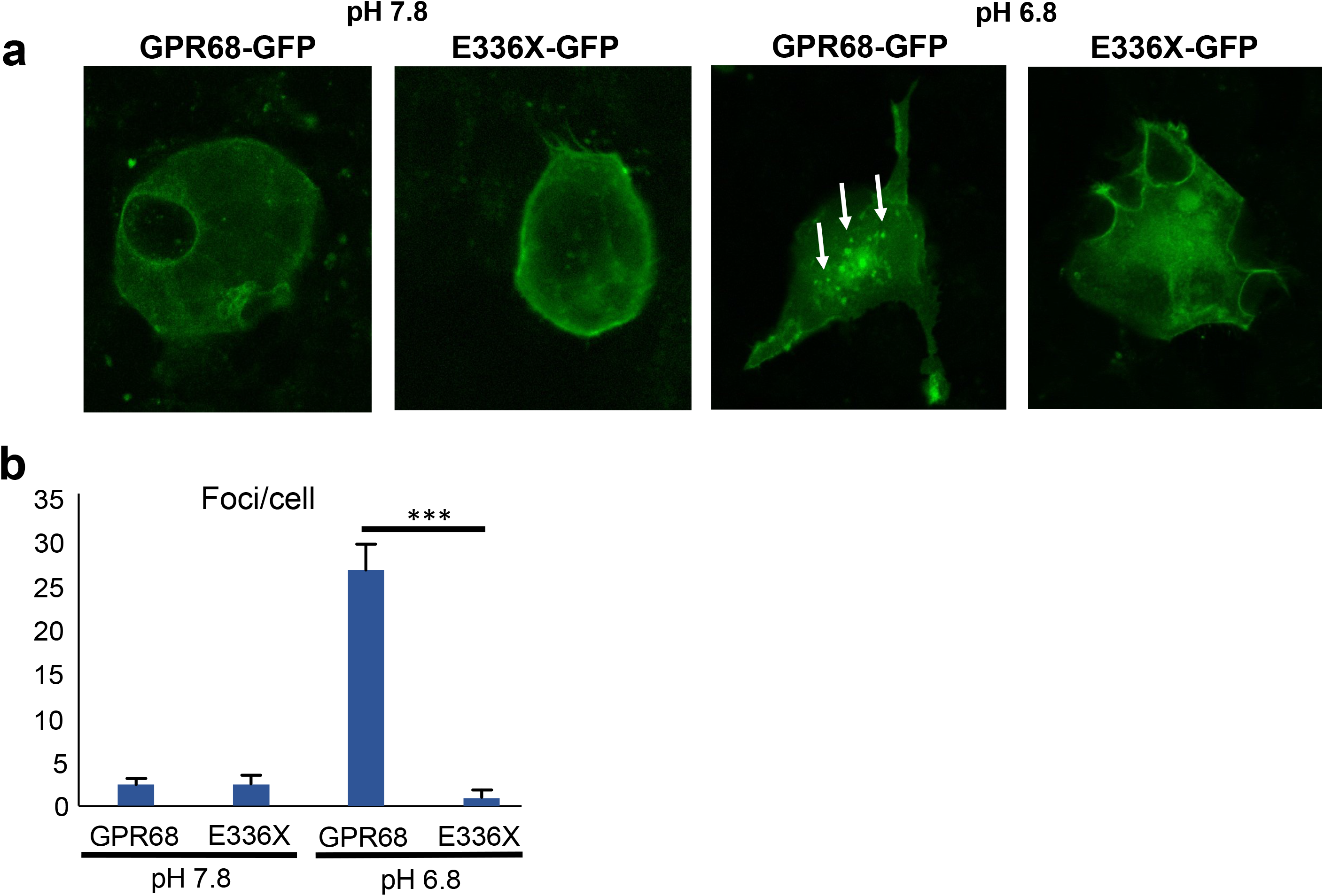
E336X does not internalize after acidification. **(a)** GPR68-GFP and E336X-GFP were transfected and had even expression through the cell membrane in pH 7.8 media. GPR68-GFP–transfected cells display punctate fluorescence, whereas E336X-GFP–transfected cells do not in pH 6.8 stimulus media. **(b)** Quantification of number of puncta per cell, n = 20.

**Supplementary Figure 11.**
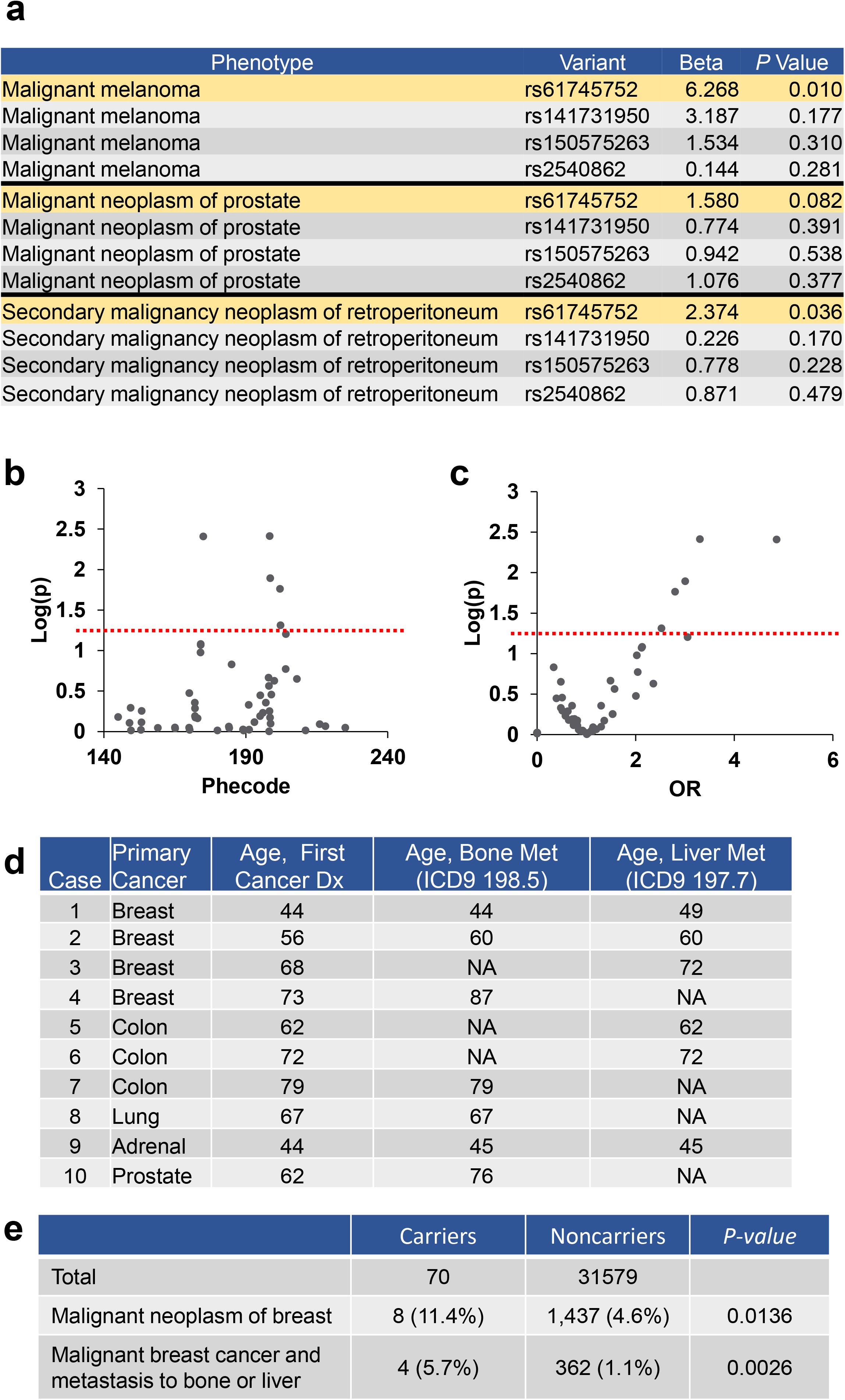
GPR68 variant Rs61745752 is associated with oncological signals including metastasis. **(a)** Analysis of the UK Biobank cohort run with a logistic regression model correcting for age, sex, and 4 principal components showed similar association with the following cancer diagnoese as the contingency analysis. Beta, beta-coefficient. MAF, minor allele frequency in UKB cohort. (**b)** Manhattan plot of targeted phenome-wide association study (PheWAS) for association of rs61745752 with cancer-related phecodes vs −log (*P*) in BioVU cohort. (**c)** Volcano plot of PheWAS for association of rs61745752 with cancer-related phecodes, plotted as odds ratio vs. −log (*P*). (**d)** Individual carriers in the BioVU with metastatic cancer, with ages at the time of cancer diagnosis and at the time of metastasis. (**e)** Comparison of the number of patients with breast cancer and the frequency of metastatic diagnosis between rs61745752 carriers and noncarriers in the BioVU cohort.

**Supplementary Figure 12.**
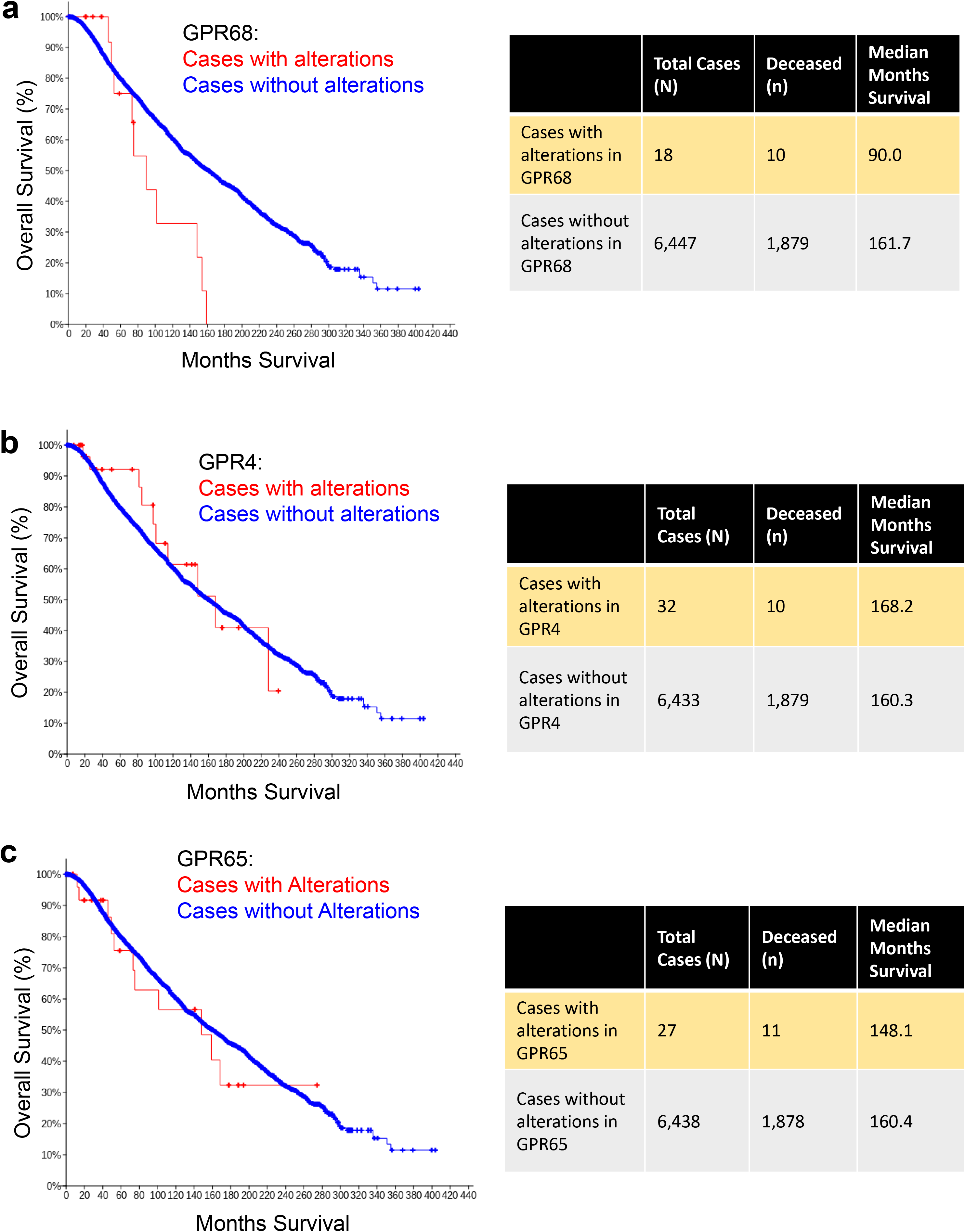
Mutations in GPR68 correlate with poor breast cancer prognosis. Kaplan-Meier plots generated in cBioportal of the breast cancer cohort carrying alterations in: (**a)** GPR68 (*P* = 0.013); (**b)** GPR4 (*P* = 0.785); (**c)** GPR65 (*P* = 0.612).

## Notes

### Competing Interest Statement

The authors have declared no competing interest.

## References

1. Vander Heiden, M. G., Cantley, L. C. & Thompson, C. B. Understanding the Warburg effect: the metabolic requirements of cell proliferation. Science 324,1029–1033 (2009).

2. Martínez-Zaguilán, R., Martinez, G. M., Gomez, A., Hendrix, M. J. & Gillies, R. J. Distinct regulation of pHin and [Ca2+] in in human melanoma cells with different metastatic potential. J. Cell Physiol. 176,196–205 (1998).

3. Sennoune, S. R. et al. Vacuolar H+-ATPase in human breast cancer cells with distinct metastatic potential: distribution and functional activity. Am. J. Physiol. Cell Physiol. 286, C1443–52 (2004).

4. Martinez-Zaguilan, R., Lynch, R. M., Martinez, G. M. & Gillies, R. J. Vacuolar-type H(+)-ATPases are functionally expressed in plasma membranes of human tumor cells. Am. J. Physiol. 265, C1015–29 (1993).

5. McLean, L. A., Roscoe, J., Jorgensen, N. K., Gorin, F. A. & Cala, P. M. Malignant gliomas display altered pH regulation by NHE1 compared with nontransformed astrocytes. Am. J. Physiol. Cell Physiol. 278, C676–88 (2000).

6. Justus, C. R., Dong, L. & Yang, L. V. Acidic tumor microenvironment and pH-sensing G protein-coupled receptors. Front. Physiol. 4, 354 (2013).

7. Webb, B. A., Chimenti, M., Jacobson, M. P. & Barber, D. L. Dysregulated pH: a perfect storm for cancer progression. Nat. Rev. Cancer 11,671–677 (2011).

8. Matthews, H., Ranson, M. & Kelso, M. J. Anti-tumour/metastasis effects of the potassium-sparing diuretic amiloride: an orally active anti-cancer drug waiting for its call-of-duty? Int. J. Cancer 129,2051–2061 (2011).

9. Seuwen, K., Ludwig, M.G. & Wolf, R. M. Receptors for protons or lipid messengers or both? J Recept Signal Transduct Res 26,599–610 (2006).

10. Ludwig, M.G. et al. Proton-sensing G-protein-coupled receptors. Nature 425,93–98 (2003).

11. Wei, W.C. et al. Coincidence Detection of Membrane Stretch and Extracellular pH by the Proton-Sensing Receptor OGR1 (GPR68). Curr. Biol. 28, 3815–3823.e4 (2018).

12. Xu, J. et al. GPR68 senses flow and is essential for vascular physiology. Cell 173, 762–775.e16 (2018).

13. Khan, M. Z. & He, L. Neuro-psychopharmacological perspective of Orphan receptors of Rhodopsin (class A) family of G protein-coupled receptors. Psychopharmacology (Berl) 234,1181–1207 (2017).

14. de Vallière, C. et al. G Protein-coupled pH-sensing Receptor OGR1 Is a Regulator of Intestinal Inflammation. Inflamm. Bowel Dis. 21,1269–1281 (2015).

15. Aoki, H., Mogi, C. & Okajima, F. Ionotropic and metabotropic proton-sensing receptors involved in airway inflammation in allergic asthma. Mediators Inflamm. 2014, 712962 (2014).

16. Okajima, F. Regulation of inflammation by extracellular acidification and proton-sensing GPCRs. Cell Signal. 25,2263–2271 (2013).

17. Wiley, S. Z., Sriram, K., Salmerón, C. & Insel, P. A. GPR68: an emerging drug target in cancer. Int. J. Mol. Sci. 20, 559 (2019).

18. Williams, C. H. et al. An in vivo chemical genetic screen identifies phosphodiesterase 4 as a pharmacological target for hedgehog signaling inhibition. Cell Rep. 11,43–50 (2015).

19. Hao, J. et al. Selective small molecule targeting β-catenin function discovered by in vivo chemical genetic screen. Cell Rep. 4,898–904 (2013).

20. Kwak, J. et al. Live image profiling of neural crest lineages in zebrafish transgenic lines. Mol. Cells 35,255–260 (2013).

21. Schilling, T. F. Genetic analysis of craniofacial development in the vertebrate embryo. Bioessays 19,459–468 (1997).

22. Birukova, A. A., Rios, A. & Birukov, K. G. Long-term cyclic stretch controls pulmonary endothelial permeability at translational and post-translational levels. Exp. Cell Res. 314,3466–3477 (2008).

23. Tian, Y., Gawlak, G., O’Donnell, J. J., Mambetsariev, I. & Birukova, A. A. Modulation of endothelial inflammation by low and high magnitude cyclic stretch. PLoS One 11, e0153387 (2016).

24. Ke, Y. et al. Mechanosensitive Rap1 activation promotes barrier function of lung vascular endothelium under cyclic stretch. Mol. Biol. Cell 30,959–974 (2019).

25. Abiko, H. et al. Rho guanine nucleotide exchange factors involved in cyclic-stretch-induced reorientation of vascular endothelial cells. J. Cell Sci. 128,1683–1695 (2015).

26. Schriek, G., Oppitz, M., Busch, C., Just, L. & Drews, U. Human SK-Mel 28 melanoma cells resume neural crest cell migration after transplantation into the chick embryo. Melanoma Res 15,225–234 (2005).

27. Sinnberg, T. et al. Wnt-signaling enhances neural crest migration of melanoma cells and induces an invasive phenotype. Mol. Cancer 17, 59 (2018).

28. Oppitz, M. et al. Non-malignant migration of B16 mouse melanoma cells in the neural crest and invasive growth in the eye cup of the chick embryo. Melanoma Res 17,17–30 (2007).

29. Schneider, L. et al. The Na+/H+ exchanger NHE1 is required for directional migration stimulated via PDGFR-alpha in the primary cilium. J. Cell Biol. 185,163–176 (2009).

30. Stüwe, L. et al. pH dependence of melanoma cell migration: protons extruded by NHE1 dominate protons of the bulk solution. J. Physiol. (Lond.) 585,351–360 (2007).

31. Stock, C. et al. pH nanoenvironment at the surface of single melanoma cells. Cell Physiol. Biochem. 20,679–686 (2007).

32. Ludwig, F. T., Schwab, A. & Stock, C. The Na+/H+ -exchanger (NHE1) generates pH nanodomains at focal adhesions. J. Cell Physiol. 228,1351–1358 (2013).

33. Stock, C. et al. Migration of human melanoma cells depends on extracellular pH and Na+/H+ exchange. J. Physiol. 567,225–238 (2005).

34. Chauhan, S. C. et al. MUC13 mucin augments pancreatic tumorigenesis. Mol. Cancer Ther. 11,24–33 (2012).

35. Su P., et al. Methods of studying mammalian cell migration and invasion in vitro. in 2017 14th International Bhurban Conference on Applied Sciences and Technology (IBCAST) 148–159 (IEEE, 2017). doi:10.1109/IBCAST.2017.7868048

36. Lourenco, S. et al. Macrophage migration inhibitory factor-CXCR4 is the dominant chemotactic axis in human mesenchymal stem cell recruitment to tumors. J. Immunol. 194,3463–3474 (2015).

37. Grinstein, S. et al. Focal localization of the NHE-1 isoform of the Na+/H+ antiport: assessment of effects on intracellular pH. EMBO J. 12,5209–5218 (1993).

38. Glunde, K. et al. Extracellular acidification alters lysosomal trafficking in human breast cancer cells. Neoplasia 5,533–545 (2003).

39. Russell, J. L. et al. Regulated expression of pH sensing G Protein-coupled receptor-68 identified through chemical biology defines a new drug target for ischemic heart disease. ACS Chem. Biol. 7,1077–1083 (2012).

40. Grover, C. N. et al. Crosslinking and composition influence the surface properties, mechanical stiffness and cell reactivity of collagen-based films. Acta Biomater. 8,3080–3090 (2012).

41. Ngo, P., Ramalingam, P., Phillips, J. A. & Furuta, G. T. Collagen gel contraction assay. Methods Mol. Biol. 341,103–109 (2006).

42. Bycroft, C. et al. The UK Biobank resource with deep phenotyping and genomic data. Nature 562,203–209 (2018).

43. Li, G. et al. Identification and characterization of distinct C-terminal domains of the human hydroxycarboxylic acid receptor-2 that are essential for receptor export, constitutive activity, desensitization, and internalization. Mol. Pharmacol. 82,1150–1161 (2012).

44. Zhou, X. E. et al. Identification of Phosphorylation Codes for Arrestin Recruitment by G Protein-Coupled Receptors. Cell 170, 457–469.e13 (2017).

45. Sente, A. et al. Molecular mechanism of modulating arrestin conformation by GPCR phosphorylation. Nat. Struct. Mol. Biol. 25,538–545 (2018).

46. Luttrell, L. M. & Lefkowitz, R. J. The role of beta-arrestins in the termination and transduction of G-protein-coupled receptor signals. J. Cell Sci. 115,455–465 (2002).

47. Ahn, S., Nelson, C. D., Garrison, T. R., Miller, W. E. & Lefkowitz, R. J. Desensitization, internalization, and signaling functions of beta-arrestins demonstrated by RNA interference. Proc. Natl. Acad. Sci. USA 100,1740–1744 (2003).

48. Carroll, R. J., Bastarache, L. & Denny, J. C. R PheWAS: data analysis and plotting tools for phenome-wide association studies in the R environment. Bioinformatics 30,2375–2376 (2014).

49. Denny, J. C. et al. Systematic comparison of phenome-wide association study of’’ electronic medical record data and genome-wide association study data. Nat. Biotechnol. 31,1102–1110 (2013).

50. Denny, J. C., Bastarache, L. & Roden, D. M. Phenome-Wide Association Studies as a Tool to Advance Precision Medicine. Annu. Rev. Genomics. Hum. Genet. 17,353–373 (2016).

51. Cerami, E. et al. The cBio cancer genomics portal: an open platform for exploring multidimensional cancer genomics data. Cancer Discov. 2,401–404 (2012).

52. Gao, J. et al. Integrative analysis of complex cancer genomics and clinical profiles using the cBioPortal. Sci. Signal. 6, pl1 (2013).

53. Gayan, S., Teli, A. & Dey, T. Inherent aggressive character of invasive and non-invasive cells dictates the in vitro migration pattern of multicellular spheroid. Sci. Rep. 7, 11527 (2017).

54. Jung, H.Y. et al. Apical-basal polarity inhibits epithelial-mesenchymal transition and tumour metastasis by PAR-complex-mediated SNAI1 degradation. Nat. Cell Biol. 21,359–371 (2019).

55. Berens, E. B., Holy, J. M., Riegel, A. T. & Wellstein, A. A cancer cell spheroid assay to assess invasion in a 3D setting. J. Vis. Exp. (2015). doi:10.3791/53409

56. Cui, X., Hartanto, Y. & Zhang, H. Advances in multicellular spheroids formation. J. R. Soc. Interface 14, 127):20160877. (2017).

57. Lin, R.Z. & Chang, H.Y. Recent advances in three-dimensional multicellular spheroid culture for biomedical research. Biotechnol. J. 3,1172–1184 (2008).

58. Mochimaru, Y. et al. Extracellular acidification activates ovarian cancer G-protein-coupled receptor 1 and GPR4 homologs of zebra fish. Biochem. Biophys. Res. Commun. 457,493–499 (2015).

59. Adams, D. S. et al. Early, H+-V-ATPase-dependent proton flux is necessary for consistent left-right patterning of non-mammalian vertebrates. Development 133,1657–1671 (2006).

60. Vandenberg, L. N., Morrie, R. D. & Adams, D. S. V-ATPase-dependent ectodermal voltage and pH regionalization are required for craniofacial morphogenesis. Dev. Dyn. 240,1889–1904 (2011).

61. Rofstad, E. K., Mathiesen, B., Kindem, K. & Galappathi, K. Acidic extracellular pH promotes experimental metastasis of human melanoma cells in athymic nude mice. Cancer Res. 66,6699–6707 (2006).

62. Nishisho, T. et al. The a3 isoform vacuolar type H^+^-ATPase promotes distant metastasis in the mouse B16 melanoma cells. Mol. Cancer Res. 9,845–855 (2011).

63. Bhujwalla, Z. M. et al. The physiological environment in cancer vascularization, invasion and metastasis. Novartis Found Symp 240, 23–38; discussion 38 (2001).

64. Hill, R. P., De Jaeger, K., Jang, A. & Cairns, R. pH, hypoxia and metastasis. Novartis Found Symp 240, 154–65; discussion 165 (2001).

65. Miraglia, E. et al. Na+/H+ exchanger activity is increased in doxorubicin-resistant human colon cancer cells and its modulation modifies the sensitivity of the cells to doxorubicin. Int. J. Cancer 115,924–929 (2005).

66. Parks, S. K., Chiche, J. & Pouysségur, J. Disrupting proton dynamics and energy metabolism for cancer therapy. Nat. Rev. Cancer 13,611–623 (2013).

67. Weiß, K. T. et al. Proton-sensing G protein-coupled receptors as regulators of cell proliferation and migration during tumor growth and wound healing. Exp. Dermatol. 26,127–132 (2017).

68. Wiley, S. Z. et al. GPR68, a proton-sensing GPCR, mediates interaction of cancer-associated fibroblasts and cancer cells. FASEB J. 32,1170–1183 (2018).

69. Castellone, R. D., Leffler, N. R., Dong, L. & Yang, L. V. Inhibition of tumor cell migration and metastasis by the proton-sensing GPR4 receptor. Cancer Lett. 312,197–208 (2011).

70. Martínez-Zaguilán, R. et al. Acidic pH enhances the invasive behavior of human melanoma cells. Clin Exp Metastasis 14,176–186 (1996).

71. Peppicelli, S., Bianchini, F., Torre, E. & Calorini, L. Contribution of acidic melanoma cells undergoing epithelial-to-mesenchymal transition to aggressiveness of non-acidic melanoma cells. Clin Exp Metastasis 31,423–433 (2014).

72. Singh, L. S. et al. Ovarian cancer G protein-coupled receptor 1, a new metastasis suppressor gene in prostate cancer. J. Natl. Cancer Inst. 99,1313–1327 (2007).

73. Ren, J. & Zhang, L. Effects of ovarian cancer G protein coupled receptor 1 on the proliferation, migration, and adhesion of human ovarian cancer cells. Chin. Med. J. (Engl). 124,1327–1332 (2011).

74. Wei, W.C. et al. Functional expression of calcium-permeable canonical transient receptor potential 4-containing channels promotes migration of medulloblastoma cells. J. Physiol. 595,5525–5544 (2017).

75. Zhu, H. et al. Proton-sensing GPCR-YAP Signalling Promotes Cancer-associated Fibroblast Activation of Mesenchymal Stem Cells. Int. J. Biol. Sci. 12,389–396 (2016).

76. Riemann, A. et al. Acidic environment activates inflammatory programs in fibroblasts via a cAMP-MAPK pathway. Biochim. Biophys. Acta 1853,299–307 (2015).

77. Horman, S. R. et al. Functional profiling of microtumors to identify cancer associated fibroblast-derived drug targets. Oncotarget 8,99913–99930 (2017).

78. Matus-Leibovitch, N., Nussenzveig, D. R., Gershengorn, M. C. & Oron, Y. Truncation of the thyrotropin-releasing hormone receptor carboxyl tail causes constitutive activity and leads to impaired responsiveness in Xenopus oocytes and AtT20 cells. J. Biol. Chem. 270,1041–1047 (1995).

79. Chen, B. et al. Small molecule-mediated disruption of Wnt-dependent signaling in tissue regeneration and cancer. Nat. Chem. Biol. 5,100–107 (2009).

80. Yu, P. B. et al. Dorsomorphin inhibits BMP signals required for embryogenesis and iron metabolism. Nat. Chem. Biol. 4,33–41 (2008).

81. Westerfield, M. The zebrafish book◻: a guide for the laboratory use of zebrafish. (University of Oregon Press, 2000).

82. Stewart, R. A. et al. Zebrafish foxd3 is selectively required for neural crest specification, migration and survival. Dev. Biol. 292,174–188 (2006).

83. Larntz, K. Small-Sample Comparisons of Exact Levels for Chi-Squared Goodness-of-Fit Statistics. J. Am. Stat. Assoc. 73, 253–263 (1978).

84. Parshall, C. G. & Kromrey, J. D. Tests of Independence in Contingency Tables with Small Samples: A Comparison of Statistical Power. Educ Psychol Meas 56,26–44 (1996).

